# Diversity, phylogeny and DNA barcoding of brachyuran crabs in artificially created mangrove environments

**DOI:** 10.1101/2020.09.07.286823

**Authors:** Ganesh Manikantan, Chinnamani PrasannaKumar, J. Vijaylaxmi, S. R. Pugazhvendan, Narra Prasanthi

## Abstract

Globally, at the rate of 1-2 percent per annum, mangrove coverings are disappearing and 35 percent have been lost in the last 20 years due to changes in climate and human activities. No mangrove-associated crabs were found when the mangroves were artificially transplanted 25 years ago in the Vellar estuary. This mangrove ecosystem was sampled for brachyuran biodiversity estimation, species abundance, composition and evaluation of the effectiveness of DNA barcoding in brachyuran crabs species identification. A total of 2844 crabs were collected, representing 35 species within 8 families belonging to 20 genera. Four brachyuran crab species, that is, *Uca lactae, U. Triangularis, Selatium brockii*, and *Neosarmatium asiaticum* account for >70% of the total abundance. An approximate 87.5% of crab species estimated to occur by various species estimator were recovered in the present study. Between *Uca lactea* and *U. triangularis*, the maximum association index value was observed (97.7%). Cluster analysis grouped the sampled stations according to the types of mangrove species, clearly influencing the structure and composition of the brachyuran crabs. In general, vegetative cover composed of multiple species of mangroves is preferred for the abundance of all collected crabs species, and particularly *Neosarmatium asiaticum*. Analysis of DNA barcoding indicates that 40% of the brachyuran species gathered in this sample were first barcoded. The advent of new high-throughput sequencing technologies will change biomonitoring applications and surveys drastically in the near future, making reference datasets like ours relevant.

## 1. Introduction

Mangrove ecosystems are of ecological and economic significance (Lee et al., 2014). With high biomass and economic values (Alongi, 2015), globally, mangroves cover 15,000,000□ha (Giri, 2011). For a variety of terrestrial and marine organisms, including several commercial species and juvenile reef fish, these forests provide food, breeding grounds and nursery sites at the land-sea interface (FAO, 2007; Igulu et al., 2014). Mangrove forests are highly productive ecosystems whose primary production rates equals those of evergreen tropical forests (Alongi, 2014). They sequester carbon in tree biomass, and by decomposition and export, much of the carbon is lost to neighbouring ecosystems (Algoni, 2012). Mangroves also play also a key role in human survival and livelihoods by being extensively useful as food, timber, fuel and medicine (Alongi, 2002; Saenger, 2003). They guard against destructive events such as tsunamis, tropical cyclones and tidal bores, and may limit shoreline erosion (Alongi et a., 2014; Igulu et al., 2014). Mangroves are disappearing at a global loss rate of 1-2 percent each year despite their importance (Spalding et al., 2010), and over the last 20 years the loss rate has reached 35 percent (FAO, 2007; Polidoro et al., 2010). Changes in climate (increasing sea level and altering rainfall events) and human activities (urbanization, aquaculture, mining, and overexploitation of timber, fish, shellfish and crustaceans) pose significant threats to mangrove ecosystems (Hsiang, 2000; McLeod et al., 2006, Gilman et al., 2008, Van Lavieren et al., 2012, Ellison et al., 2012, Carugati et al., 2018; Elwin et al., 2019).

Anthropogenically contaminated mangrove area reported a 20% loss of benthic biodiversity, with the local extinction of major macrobenthic phyla, a loss of 80% of microbial-mediated rates of decomposition, benthic biomass and trophic resources (Carugati et al., 2018). Unlike naturally formed mangrove forests (like Tampa Bay, Florida), where tidal salt marshes were naturally transposed into mangrove wetlands (Osland et al., 2012), Vellar estuary mangroves were planned and risen from mangrove saplings (see, Ajmal Khan et al., 2005; Kathiresan & Rajendran, 2005). One of the fertile estuaries in Tamil Nadu is the Vellar estuary at Parangipettai (lat. 11°29’N; long. 79°46’E), which flows over the southeast coast of India.. In 1991, nearly 1.5 km upstream of the mouth in the tidal zone and on the northern shore of the estuary, a mangrove plantation covering an area of 10 ha was created (Kathiresan and Rajendran, 2005; Sandilyan & Kathiresan, 2012). During 2004, Vellar estuary mangroves played a significant role in reducing major tsunami impact. Mangrove forests in the Indo-Pacific host abundant molluscs and crab species, both of which control the energy and food web of the ecosystem (Plaziat, 1984; Fratini et al., 2004; Cannicci et al., 2008; Sandilyan & Kathiresan, 2012). Brachuyuran crabs are as numerous as molluscs in terms of diversity (Cannicci et al., 2008). For e.g., in the Indian subcontinent, 149 brachyuran crab species belonging to 75 genera have been found living in mangroves (Dev Roy, 2008), with > 100 species known to colonise peninsular Malaysian mangroves (Tan and Pg, 1994).

Mitochondrial DNA exhibits many features in most animals that make it highly desirable for the molecular identification, like occurring in a high number of mitochondrial copies per cell, almost entirely maternal inheritance, the absence of introns and recombination (Ballard and Whitlock, 2004; Ballard and Rand, 2005; Bernt et al., 2013). The standardised use of sequencing and archiving of an approximately 650 base pair (bp) fragment of the mitochondrial cytochrome c oxidase subunit 1 (COI) represents a very effective DNA-based method (DNA barcoding), for the identification of animal specimens (Hebert et al., 2003a; 2003b; Bucklin et al., 2011). DNA barcoding’s two main objectives are (i) to assign unknown specimens to species that have already been described and identified, and (ii) to improve the discovery of new species (including cryptic, microscopic, and organisms with inaccessible or complex morphological characteristics) (Hebert et al., 2003a, b). As a result, some recently published species descriptions and taxonomic studies (including immature life stages of brachyurans) have included barcode sequence data (Markert et al., 2005; Chen et al., 2012; Riehl and Kaiser, 2012; PrasannaKumar et al., 2012; De Grave et al., 2013; Meiklejohn et al., 2013; Khalaji-Pirbalouty and Raupach, 2014; Feroz et al., 2014; Shin et al., 2014; Gunalan and Prasannakumar, 2014; Brandão et al., 2016; Rahman et al., 2019; Egli et al., 2020) and several studies have been published for the pioneer efforts in barcoding a particular or a group of species including barchyurans (Kelnarova et al., 2018; Brown, 2019; Biella et al., 2020; Madhavan et al., 2020; PrasannaKumar et al., 2020a). The usefulness of DNA barcoding is now well established for biodiversity and conservation (von Cräutlein et al., 2011; Krishnamurthy and Francis, 2012; Li et al., 2017; Beng and Corlett, 2020). Several previous studies have used DNA barcoding to test the biodiversity of different marine flora (including mangroves) and fauna (besides brachyurans) in these artificially transplanted mangrove-rich Vellar estuary (Khan et al., 2010, 2011; Rahman et al., 2013; PrasannaKumar et al., 2012; Thirumaraiselvi et al., 2015; Rajthilak et al., 2015; Sahu et al., 2016; Hemalatha et al., 2016; Palanisamy et al., 2020; PrasannaKumar et al., 2020a; 2020b; 2020c; Manikantan et al., 2020; Thangaraj et al., 2020; Prasanthi et al., 2020).

There were no mangrove-associated brachyuran crabs when the mangroves were artificially transplanted into the Vellar estuary (Kathiresen and Rajendran, 2005; Ajmal Khan et al., 2005). Parallel to the transplanted Vellar mangroves, the Pichavaram mangrove ecosystem (11.4319 ° N, 79.7810 ° E) is a heterogeneous mixture of mangrove elements distributed over 10 km^2^ and host about 38 species of brachyuran crabs (Ajmal Khan et al., 2005). A previous study sampled 8 brachyuran crab species in the Vellar estuary (Ajmal Khan et al., 2005) and speculated that the remaining species of Pitchavaram mangrove will also colonise the transplanted Vellar mangroves in due course of time, but how long was then uncertain. The growth of mangroves influences the biogeochemistry of the soil. For example, after continuous influencing soil biogeochemistry for 11 years, *Rhizophora mucronata* saplings took approximately 25 years to reach full maturity in Philippines (Salmo et al., 2013). We aim to check the biodiversity of brachyuran crab species in Vellar mangroves, since it’s been nearly 25 years after sampling transplantation. In this study, we intent to estimate the diversity, species richness, composition and phylogenetic relationships of barchyuran crab species inhabiting in artificially created habitats of Vellar mangrove. We also aim to evaluate the efficacy of DNA barcoding technique in precisely identifying brachyuran crab species of Vellar mangroves.

## 2. Materials and methods

### 2.1. Sample collection and identification

Between February 2015 and 2016, in Southeast coast of India (**Fig. S1**), extensive collection was carried out on mangrove patches along the shores of the Vellar estuary at Parangipettai ((lat. 11°29’N; long. 79°46’E). *Rhizophora* spp., *Avicennia* spp., and *Acanthus illicifolius* were major species distributed along the estuary shore as single and mixed patches of vegetation. Eight species of brachyuran crabs with a density of 27-40/m^2^ have been reported a decade after their establishment (Ajmal Khan et al., 2005). The present study samples brachyuran crabs along the estuary shore at 10 sampling stations. Based on the mangrove species composition at each station, the sampled stations were classified into 4 types; *viz*., 1. Mixed mangrove patch (M) (containing *Avicennia marina, A. officinalis, Rhizophora apiculata, R. mucronata* and *R. annamalayana)*, 2. *Avicennia marina* patch (A) (containing *Avicennia marina*), 3. *Acanthus ilicifolius* patch (Ai) (containing *Acanthus ilicifolius*) and 4. Non-mangrove patch (N) dominated by the salt marshes (containing *Suaeda maritima* and *Prosopis juliflora)*. To estimate crab density, a random sampling approach (using a randomly placed low tide 5 square metre quadrate) was used (Ajmal Khan et al., 2005). Crabs were washed in ambient waters, transported to the laboratory, where they were maintained until further study in −20°C. Based on the keys given by Sakai (1976), Sethuramalingam and Ajmal Khan (1991), Ragionieri et al. (2012) and Lee et al. (2015), crab species were identified.

### 2.2. Abundance and diversity estimation

An approximation of the abundance and diversity was made using Plymouth Routines in Multivariate Ecological Research (PRIMER) version 6.1.10 (Clarke and Warwick, 2001). To compare species diversity between stations, the dominance plot (Warwick, (1986), which involves the plotting of different k-dominance curves (Lambshed et al., 1983) on the X-axis (logarithmic scale) with percent dominance on the Y-axis (cumulative scale), was used.

To get more detailed data on bracyuran crab diversity, we deployed a variety of diversity indices *viz*., Margalef index (based on the number of species at a site; -species richness), the Pielou’s evenness index (calculates evenness based on the observed diversity; Pielou, 1969), the Brillouin alpha index (based on total number of individuals in the samples and number of individuals per species), the Fisher alpha index (based on the assumption that species abundance follows the log series distribution), rarefaction index (predicts species richness based on the number of samples), Shannon – Wiener index (based on the proportion of individuals in the individual species), Simpson’s index (based on the number of individuals per species and the total number of the individuals in the sample), taxonomic diversity index (based on the taxonomic relation between different organisms in a community), taxonomic distinctness index (based on the taxonomic relatedness and evenness), total phylogenetic diversity (based on phylogeny).

Similarly, to obtain estimates of the total number of species likely to occur with intensive selection in the study field, the following estimators were used: Sobs (based on the total number of species found in a sample or sample sets), Chao 1 (based on the approximate true species diversity of a sample) and Chao 2 (based on the presence-absence data of a species in the sample), Jacknife 1 (based on the number of samples and the number of species in any one sample), Jacknife 2 (based on the number of species only found in one sample and number of species only found in two samples), Bootstrap (based on the presence of individual species in the proportion of sample randomly resampled), Michaelis Menton (based on the species diversity and richness) and Ugland-Gray-Ellingson (based on how many taxa would have been found at each site if a specific number of sampling units [=□quadrats] had been sampled at each site). Using the association index (Smith et al., 1993) in PRIMER ver. 6.1.10., the calculation of the degree of linkage equilibrium within the collected species, sampled stations and influences of the mangrove types were calculated.

### 2.3. Statistical analysis

The multivariate approaches, unlike diversity indices, maintain species identity and are more susceptible to detecting population trends and thus detecting subsequent effects (Warwick and Clarke, 1991). Cluster analysis was used to cluster associated groups by hierarchical agglomerative methods using the Bray-Curtis coefficient (Bray and Curtis, 1957) to produce the dendrogram. To compare the diversity between sampled stations, the cophenetic correlation coefficient was used. In order to construct a ‘map’ to demonstrate the interrelationships between all entities, non-metric Multi Dimensional Scaling (MDS) was used to determine the similarities (or dissimilarities) between each pair of entities. To get a sense of how well the two-dimensional data is organised and matches the original data patterns, Shepard’s plot was drawn. To expose associations between mangrove and crab species forms, the bubble plot was synthesised. In order to quantify the contribution (percent) of each species to the dissimilarity values between types of mangroves in the sampled stations, SIMPER analysis was performed. To test the respective sample’s powerful discriminators, SIMPER uses Bray-Curtis dissimilarity.

### 2.4. DNA barcoding and analysis

For DNA barcoding, 15 brachyuran crab species that were morphologically difficult to differentiate were selected, *viz., 1-Neosarmatium asiaticum, 2-Episesarma versicolor, 3-Perisesarma bidens, 4-Parasesarma plicatum, 5-Nanosesarma minutum, 6-Plagusia dentipes, 7-Ocypode platytarsis, 8-Cardisoma carnifex, 9-Macrophthalmus depressus, 10-Metopograpsus frontalis, 11-Metopograpsus latifrons, 12-Grapsus albolineatus, 13-Uca lacteal, 14-Uca triangularis and 15-Ocypode brevicornis*.

Approximately 50-100 mg of tissue (from the second pereopod) was used for DNA extraction using the NucleoSpin® Tissue Kit (Macherey-Nagel) following the instructions of the manufacturer. The tissues with lysis buffer and proteinase K were incubated at 56°C until complete tissue lysis, after which 5 μl of RNase A (100 mg/ml) was added and incubated at room temperature for 5 minutes. Using spin columns supplied with the kit, DNA was eluted and stored at −20 °C until further analysis. The eluted DNA in the 50 μl buffer was used as such (without dilution) for the polymerase chain reaction (PCR). Folmer’s primer pairs; LCO1490 (5’-GGTCAACAAATCATAAAGATATTGG-3’) and HCO2198 (5’-TAAACTTCAGGGTGACCAAAAAATCA-3’) (Folmer et al., 1994) were used to amplify the COI gene fragments using PCR conditions; initial denaturation at 98°C (30 seconds), 10 cycles of 98°C (5 seconds), 45°C (10 seconds), 72°C (15 seconds), followed by 30 cycles of 98°C (5 seconds), 50°C (10 seconds), 72°C (15 seconds) and the final extension at 72°C (60 seconds). Sangar’s dideoxy sequencing (two ways) was performed using DNA Analyzer 3730xL (Applied Biosystems, USA) in Rajiv Gandhi Centre for Biotechnology, Trivandrum (India).

Using Sequence Scanner Software v1 (Applied Biosystems) and Geneious Pro v5.1 (Drummond et al., 2010), sequence quality and the alignment was respectively carried out.

Basic Local Alignment Search Tool (BLAST) algorithms (Altschul et al., 1997) were used for identifying DNA sequences by comparing with the GenBank database. Based on 2% cut-off values for phylogram tree based identification, the Genbank reference sequences were selected. If there are 2 or more sequences with equivalent similarity values (of the same species), only one sequence is selected on the basis of the highest query coverage (q) or the lowest error value (e). If 2 or more sequences with equal similarity values, and exact q and e values occurs, the first sequences was selected. Neighbor-Joining trees were constructed and pair-wise distance was calculated using Kimura-2 parametric (K2P) distance model (Kimura, 1980) using 1000 bootstrapping (Nei and Kumar, 2000).

## 3. Results

### 3.1. Collection, classification and diversity estimation

Station 1 was known for *Avicennia marina* patch (and indicated as 1A) based on the presence of mangrove species; Station 2 was non-mangrove zone containing *Suaeda maritima* and *Prosopis juliflora* (2N); station 3 was mixed mangrove zone (3M) (containing 5 mangrove species; *viz., Avicennia marina, A. officinalis, Rhizophora apiculata, R. mucronata* and *R. annamalayana*); Station 4 had single species of *Avicennia marina* (4A); Station 5 was mixed mangrove (5M) types (composition similar to station 3M excerpt the absence of *R. mucronata* and *R. annamalayana*); Station 6 had single species of *Acanthus ilicifolius* (6Ai); Station 7 represented non-mangrove zone (7N) (similar of 2N); Station 8 had single species of *Avicennia officinalis* (8A); Station 9 had single species of *Acanthus ilicifolius* (9Ai) and Station 10 had single species of *Avicennia marina* (10A).

A total of 2844 crabs were collected, representing 35 species (Table S1). These 35 species belonged to 20 genera within 8 families (Sesarmidae, Portunidae, Ocypodidae, Grapsidae, Dotillidae, Varunidae, Macrophthalmidae and Gecarcinidae). Barchyuran crabs were richer in station 3 (M) and meagre in station 1 (A) (**Fig. 1**). At station 3M the dominance plot revealed higher diversity as the natural curve representing station (3M) dropped at the bottom (**Fig. S2**). Also alpha diversity indices such as number of species (S); abundance (N); Margalef index (d), Brillouin, Fisher; rarefaction index (ES - 25, 50, 75, 100, 150), Shannon-Wiener diversity index (H’(log2), total taxonomic distinctness index (sDelta+) and the total phylogenetic diversity index (sPhi+) highlighted the higher diversity in mixed mangrove patch at 3M (**Fig. S2.A, Table S2**). However, the Pielou’s evenness index (J’), the Simpson dominance index (Lambda’), the Simpson richness index (1-Lambda’) showed higher diversity at 1A and highest taxonomic diversity index (Delta) at 2N (**Table S2**). But based on the shade plot and dominant plot curves, these exceptions could be rejected.

**Fig. 1:**
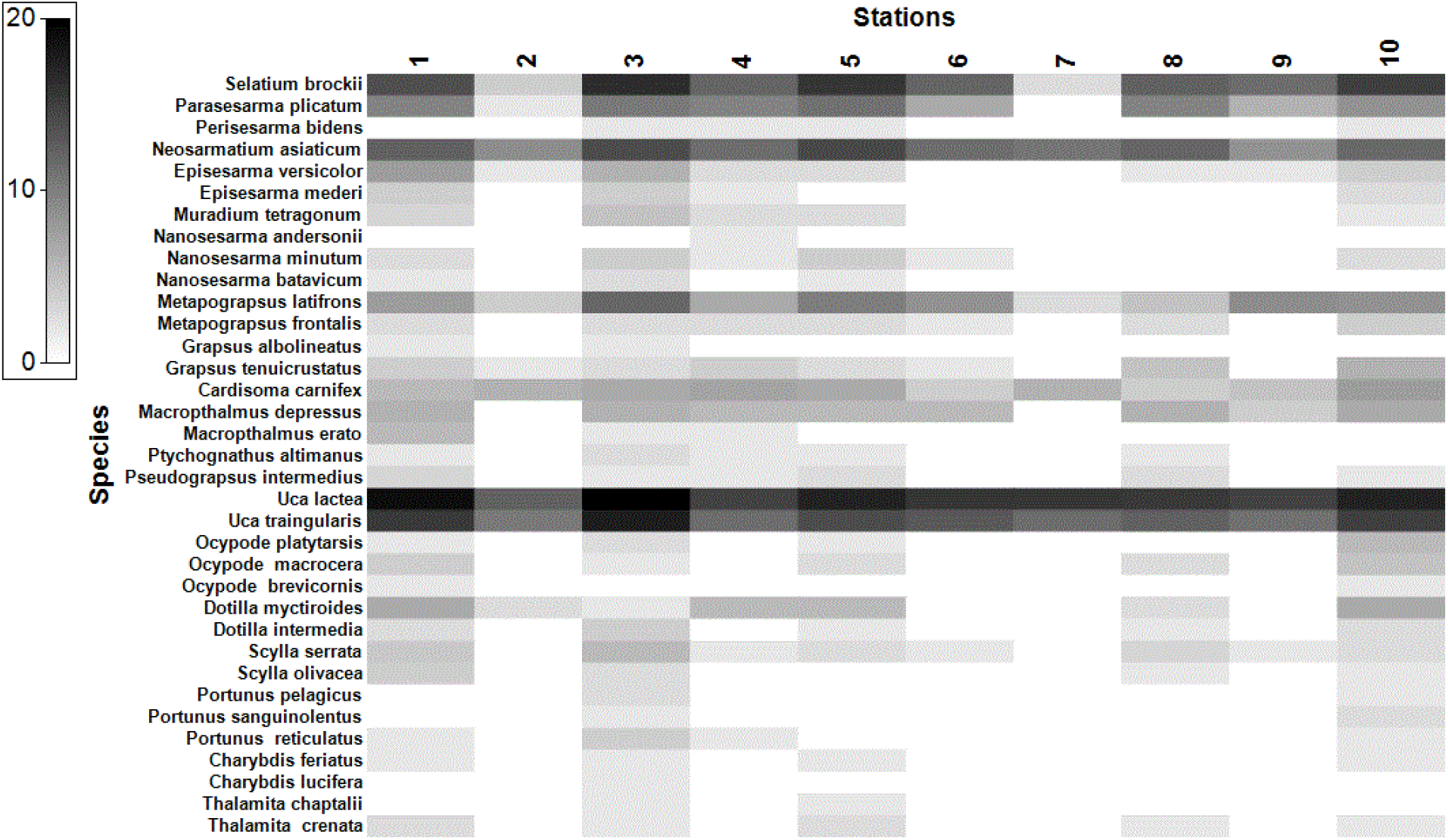
**S**hade plot represents the abundance abundance of brachyuran crab species. Note in all sampled stations, species such as *Uca lactea, Uca triangularis* and *Neosarmatium asiaticum* were consistently abundant.

The 8 families, *viz*., Sesarmidae, Portunidae, Ocypodidae, Grapsidae, Dotillidae, Varunidae, Macrophthalmidae and Gecarcinidae were represented by 10, 9, 5, 4, 2, 2, 2 and 1 species, respectively (**Table S1**). Four crab species cumulatively contributes >70% of total abundance, *viz., Uca lactae* (~26%), *U. triangularis* (17.3%), *Selatium brockii* (14.8%), and *Neosarmatium asiaticum* (13.2%) (**Table S1**). The number of brachyuran crab species varied between the minimum of 6 species at 7N and maximum 33 species at 3M.

The abundance varied from the minimum of 117 crabs at 2N and maximum of 480 crabs extracted at 3M (**Table S1**). Four sampled stations, *viz*., 3M (~17%), 1A (~14%), 5M (13.6%) and 10A (13.2%), contributed >57% of the abundance. Less than 10% abundance was contributed by each other stations. Among all the sampled stations, the estimated species (ES) numbers for 25, 50, 75,100 and 150 crabs were below 33 species. Thus the sampled sizes were found to be reasonably adequate for the number species reported (n=35) (**Table S2**). However, different species estimators, *viz*., Sobs, Chao 1, Chao 2, Jacknife 1, Jacknife 2, Bootstrap, Michaelis Menton (MM) and Ugland-Gray-Ellingsen (UGE), used to estimate the true total number of species that would have been observed with intensive sampling (assuming a closed population is sampled successively), predicted a maximum of 39.86 (by Chao 2 in 3M) (**Fig. S2B, Table S2**). With intense sampling there is a possibility to obtain 40 crab species from the ten stations. While the present study recovered 87.5% (n=35) of a total species estimated.

### 3.2. Index of association of among the sampled species and stations

The index of association analysis between the sampled species suggested the maximum association value (97.7%) between *Uca lactea* and *U. triangularis* (**Table S3**). When grouping sampled stations to determine the index of association, 9Ai and 6Ai showed the highest association value (89.8%) (**Table 1**). The minimum association value (43.1%) was recorded between the stations; 7N and 1A (**Table 1**).

**Table 1.** Results for index of association index (lower diagonal) and mangrove types (upper diagonal) for all sampled stations. In both cases, the overall association value of 9Ai and 6Ai was >88%. Higher and lower statistical values were highlighted in bold.

| Stations | 1A | 2N | 3M | 4A | 5M | 6Ai | 7N | 8A | 9Ai | 10A |
| --- | --- | --- | --- | --- | --- | --- | --- | --- | --- | --- |
| 1A |  | 49.94 | 84.64 | 78.97 | 83.77 | 68.42 | 46.74 | 78.76 | 61.83 | 85.13 |
| 2N | 58.26 |  | 46.49 | 64.51 | 55.93 | 68.14 | 83.66 | 65.06 | 74.74 | 52.47 |
| 3M | 86.13 | 57.01 |  | 73.77 | 84.25 | 65.10 | **44.15** | 71.98 | 59.11 | 81.33 |
| 4A | 79.61 | 66.62 | 78.67 |  | 82.46 | 80.90 | 59.63 | 82.77 | 76.05 | 79.68 |
| 5M | 82.90 | 63.12 | 83.39 | 85.36 |  | 75.53 | 52.53 | 81.89 | 69.07 | 84.17 |
| 6Ai | 64.00 | 72.17 | 66.69 | 74.89 | 75.86 |  | 70.64 | 82.10 | **88.96** | 71.46 |
| 7N | **43.00** | 81.11 | 44.37 | 50.54 | 48.55 | 63.12 |  | 60.87 | 72.61 | 49.20 |
| 8A | 77.79 | 64.74 | 73.91 | 80.66 | 82.90 | 78.37 | 51.63 |  | 76.43 | 78.08 |
| 9Ai | 61.29 | 73.77 | 64.36 | 73.22 | 72.05 | **89.89** | 64.37 | 75.85 |  | 64.79 |
| 10A | 85.10 | 60.83 | 84.02 | 82.31 | 83.24 | 67.29 | 45.06 | 77.94 | 64.00 |  |

Excerpt for 6 species (*viz., Selatium brockii, Neosarmatium asiaticum, Metapograpsus latifrons, Cardisoma carnifex, Uca lactea*, and *U. triangularis*), all other species present in 1A were absent in 7N (**Table S1**). Similarly, when mangrove types were grouped on the basis of similarity 9Ai and 6Ai were very similar (88.96%). The stations 7N and 3M (44.15%) were very dissimilar (**Table 1**). Interesting to note that the sampled stations were grouped according to the mangrove types (A, Ai, M & N). However such patterns of groupings were not noticed among the brachyuran crabs species (excerpt *Uca* spp.). In this regard, further investigations could be directed to determine the factors shaping the association indices of brachyuran crabs in artificially created mangrove ecosystems.

### 3.3. Statistical analysis

The cophenetic correlation was found to be strong (0.809). All of the grouping patterns observed in the MDS plot had higher similarity levels (>80%) (**Fig. 2**). The first grouping was of 88.96% similarity between 6Ai and 9Ai, which revalidated the reliability of the aggregation of index. So, for the given mangrove species (like *Acanthus ilicifolius*) there was an established pattern of bracyuran crab community. In other words, type of mangrove species influences the structure and species composition of brachyuran crabs. Similar patterning at non-mangrove (2N and 7N) stations was also observed. However, the 4 sampled stations that contains *Avicennia* (*viz*., 1A, 4A, 8A and 10A) have been segregated into two distinct groups (**Fig. 2.A**).

**Fig. 2:**
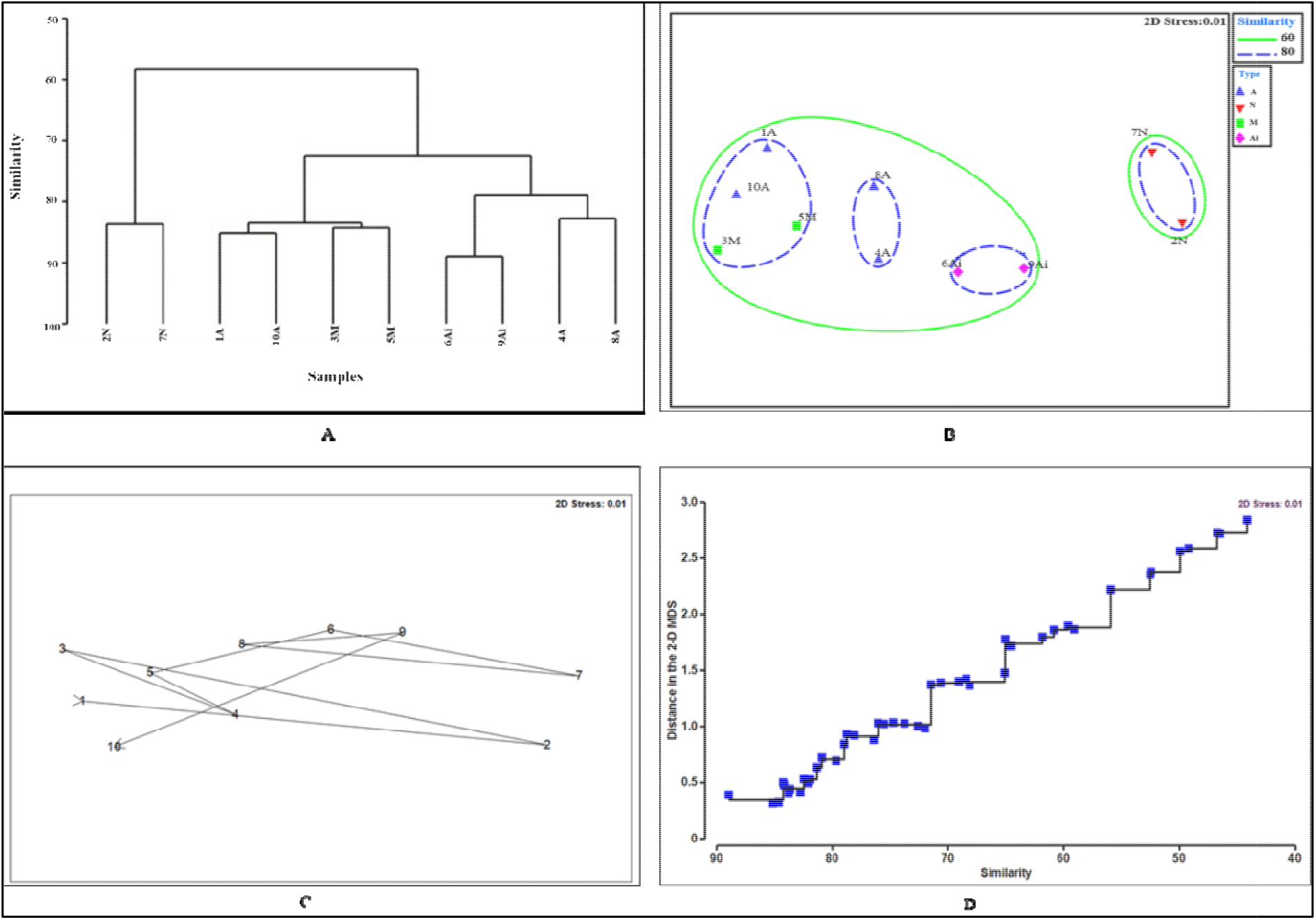
Cluster analysis results (A) are represented by a dendrogram with the x-axis representing the collections of samples, and the y-axis describing the degree of similarity at which the samples or groups are fused. Combined MDS and cluster plot (B) shows that closer lying samples were similar in species composition and vice versa. The MDS trajectory plot (C) interconnects the sampled stations where the line’s direction and length indicates similarity (or distance) between stations. Shepard plot (D) shows a good fit (stress value; 0.01) between the distances of the community (similarity) and the distances of ordination (indicated by the monotonic black regression line).

Non-metric multidimensional scaling (MDS) was plotted to account for this distinct grouping pattern of *Avicennia* stations. MDS showed that stations with mangroves (mixed species and *Avicennia*) fell to the left side of plot, stations with Acanthus *ilicifolius* (6Ai and 9Ai) fell to the middle and non-mangrove zone (2N and 7N) fell to the right side of the MDS plot (fig. 2B). Even though while clustering all stations, >80% similarity level was observed, the stations with mangroves (mixed, *Avicennia* and *Acanthus*) and non-mangroves formed two distinct clusters in the plot when the similarity level was reduced to 60%. Stations with mixed mangrove species; 5M (with dominant *Avicennia* spp. populations (*A. marina* and *A. officinalis*) and a *Rhizophora apiculata*) was positioned relatively closer to other *Avicennia* mangrove stations (6Ai and 9Ai) than mixed mangrove station 3M (dominated by 3 *Rhizopora* spp. (*Rhizophora apiculata, R. mucronata* and *R. annamalayana*) and *Avicennia* spp. (*A. marina* and *A. officinalis*)) (**Fig. S1**).

*Rhizopora* spp. richer in 3M than 5M was also noticed. So it was found that, *Avicennia* spp. dominating 5M station was the reason for distinct clustering patterns of 1A, 10A and 4A, 8A in the dendrogram. But it has been unclear how a mangrove species (in a mixed mangrove community) affect a crab species (in mixed population). This does not, however, alter the observed fact that the type of mangrove species affects the structure and species composition brachyuran crabs, because the pattern of crab species composition depends on the presence and absence of mangrove species (**Fig. 2.B**). The MDS trajectory plot (**Fig. 2.C**) also showed that the types of mangrove species (rather than its geographical distances) determines the species compositions of colonising brachyuran crabs. This trend was reconfirmed by the Shepard stress plot (**Fig. 2.D**), as a good fit of ordination was observed. The stress value was very poor (0.01), and the points dropped nearer to the line of regression line (**Fig. 2.D**).

### 3.4. Bubble segmented plot and SIMPER analysis

We used bubble segmented plot in order to understand the essence of inter-relationship between mangroves and crab population at the given sampled stations. The plots were drawn from all all sampled stations for the top 4 abundant crab species (A - *Selatium brockii*, B - *Neosarmatium asiaticum*, C - *Uca lactea* and D - *Uca triangularis*) (**Fig. 3**). When the number of *Rhizopora* spp. decreases in the stations of mixed mangrove population (5M > 3M), the abundances of *Uca lactea* and *U. triangularis* decreased. *Selatium brockii* abundances have been significantly reduced at non-mangrove stations (7N and 2N), exposing this species’ heavy dependence on mangrove vegetation (**Fig. 2**). In general, the abundances of all collected crab species and *Neosarmatium asiaticum* in particular, prefers mixed mangrove vegetation, followed by *Avicennia* spp., *Acanthus ilicifolius* and non-mangrove vegetation.

**Fig. 3:**
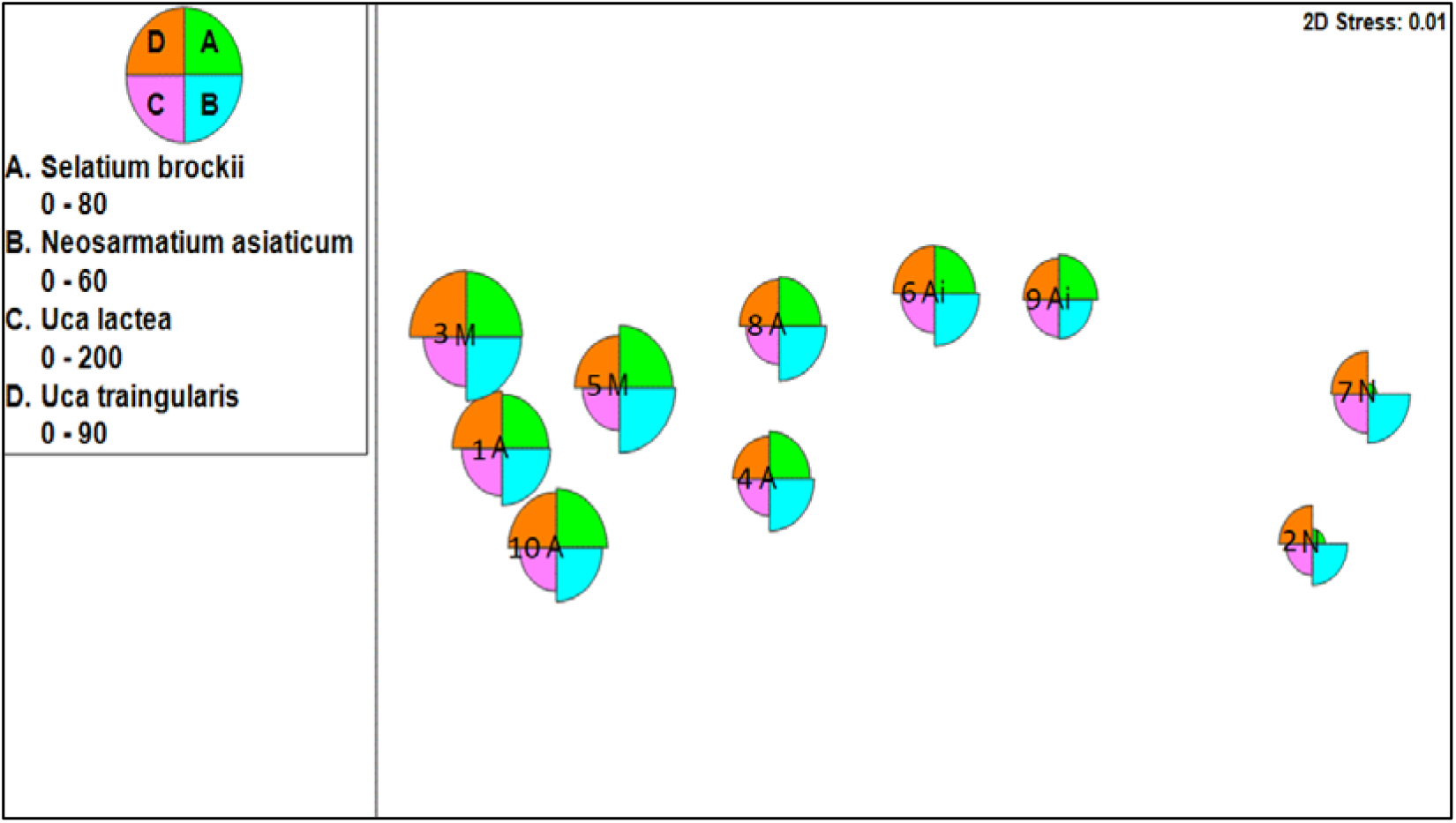
Segmented bubble plot showing distribution of top four abundant brachyuran crab species between the sampled stations. The abundances of crab species declines in the following patterns; mixed mangrove types (highest abundances and species diversity), followed by *Avicennia* spp. types, *Acanthus ilicifolius* types and non-mangrove types.

The average similarity for the sampled stations; M, A, Ai and N were 84.26%, 80.57%, 88.96% and 83.66%, respectively (**Fig. S3**). While *Uca lactea* contributed maximum average similarity for all four sampled stations types (between 11.04% in M and 23.95% in N), different species contributed minimum average similarity for each sampled mangrove types (**Fig. S2**). *Macropthalmus depressus* (3.45%), *Cardisoma carnifex* (3.7%), *Metapograpsus latifrons* (10.84%) and *Neosarmatium asiaticum* (17.33%), respectively contributed the minimum average similarity values for types M, A, Ai and N (**Fig. 3**). The factors driving the minimum contribution values for different brachyuran crab species at a given mangrove type station is currently unknown. Maximum average dissimilarity (MAD) was observed between M and N stations (50.22%) (**Table 2**). *Selatium brockii* is the only species to contribute to MAD observed between N and all types of mangrove stations.

**Table 2:** Average dissimilarity between the mangrove types was shown in percentage. Percentage values in the parenthesis represents the average percentage dissimilarity values. Values in lower and upper diagonals represents the maximum and minimum average dissimilarity percentage contribution of brachyuran crab species, respectively.

|  | M | A | Ai | N |
| --- | --- | --- | --- | --- |
| **M** |  | *Ocypode macrocera*<br>(0.56%) | *Metapograpsus latifrons* (0.94%) | *Pseudograpsus intermedius* (1.13 %) |
| **A** | 19.49%<br>*Metapograpsus latifrons* (1.47%) |  | *Ocypode platytarsis* (0.79 %) | *Neosarmatium asiaticum* (1.11%) |
| **Ai** | 32.79%<br>*Parasesarma plicatum* (1.92%) | 27.25%<br>*Dotilla myctiroides* (2.49%) |  | *Cardisoma carnifex* (1.36 %) |
| **N** | 50.22%<br>*Selatium brockii* (6.19%) | 43.94%<br>*Selatium brockii* (5.49%) | 28.46%<br>*Selatium brockii* (6.53%) |  |

### 3.5. DNA barcoding and pairwise analysis

BLAST analysis revealed that 40% of the species (*viz., Plagusia dentipes, Ocypode platytarsis, O. brevicornis, Metopograpsus frontalis*, and *M. latifrons* and the genus *Macrophthalmus* sp. (*M. depressus*)) was barcoded for the first time (**Table 3**). The maximum similarity values of the aforementioned species was <95% with that of all barchyuran COI sequences in the Genbank database, and individual searches for these species revealed their absence in the Genbank. *Macrophthalmus depressus* shared a similarity value of only 86% (with *Uca* spp.) and the COI sequences of none of the species belonging to this genera was found in the Genbank database suggesting that, the genus *Macrophthalmus* sp. was first time barcoded.

**Table 3.** Accession numbers of 15 species and the species barcoded for the first time were provided with their corresponding percentage of similarity and closest species matches.

| No. | Name of the species | Accession Number | Family | First time sequenced specimens with closest Genbank reference |
| --- | --- | --- | --- | --- |
| 1. | *Neosarmatium asiaticum* | KY242627 | Sesarmidae | - |
| 2. | *Episesarma versicolor* | KY284641 | Sesarmidae | - |
| 3. | *Perisesarma bidens* | KY436763 | Sesarmidae | - |
| 4. | *Parasesarma plicatum* | KY284646 | Sesarmidae | - |
| 5. | *Nanosesarma minutum* | KY284644 | Sesarmidae | - |
| 6. | ***Plagusia dentipes*** | KY284645 | Plagusiidae | 92% matched with *P. immaculata* |
| 7. | *Cardisoma carnifex* | KY398729 | Gecarcinidae | - |
| 8. | ***Macrophthalmus depressus*** | KY427938 | Macrophthalmidae | 86% similar with *Uca stenodactylus* |
| 9. | ***Metopograpsus frontalis*** | KY284642 | Grapsidae | 94% similar with *M. thukuhar* |
| 10. | ***Metopograpsus latifrons*** | KY284643 | Grapsidae | 94% similar with *M. thukuhar* |
| 11. | *Grapsus albolineatus* | KY436762 | Grapsidae | - |
| 12. | *Uca lactea* | KY406231 | Ocypodidae | - |
| 13. | *Uca triangularis* | KY284640 | Ocypodidae | - |
| 14. | ***Ocypode brevicornis*** | KY284647 | Ocypodidae | 89% similar with *O. platytarsis* |
| 15. | ***Ocypode platytarsis*** | KY284648 | Ocypodidae | 95% similar with *O. macrocera* |

The COI Sequences were grouped into 3 major clades (Sesarmidae-Plagusiidae-Macrophthalmidae (SPM) clade; Ocypodidae-Grapsidae (OG) clade and Gecarcinidae-Grapsidae (GG)) in the constructed phylogram (**Fig. 4**) by clearly distinguishing the out group (*Penaeus monodon*).

**Fig. 4:**
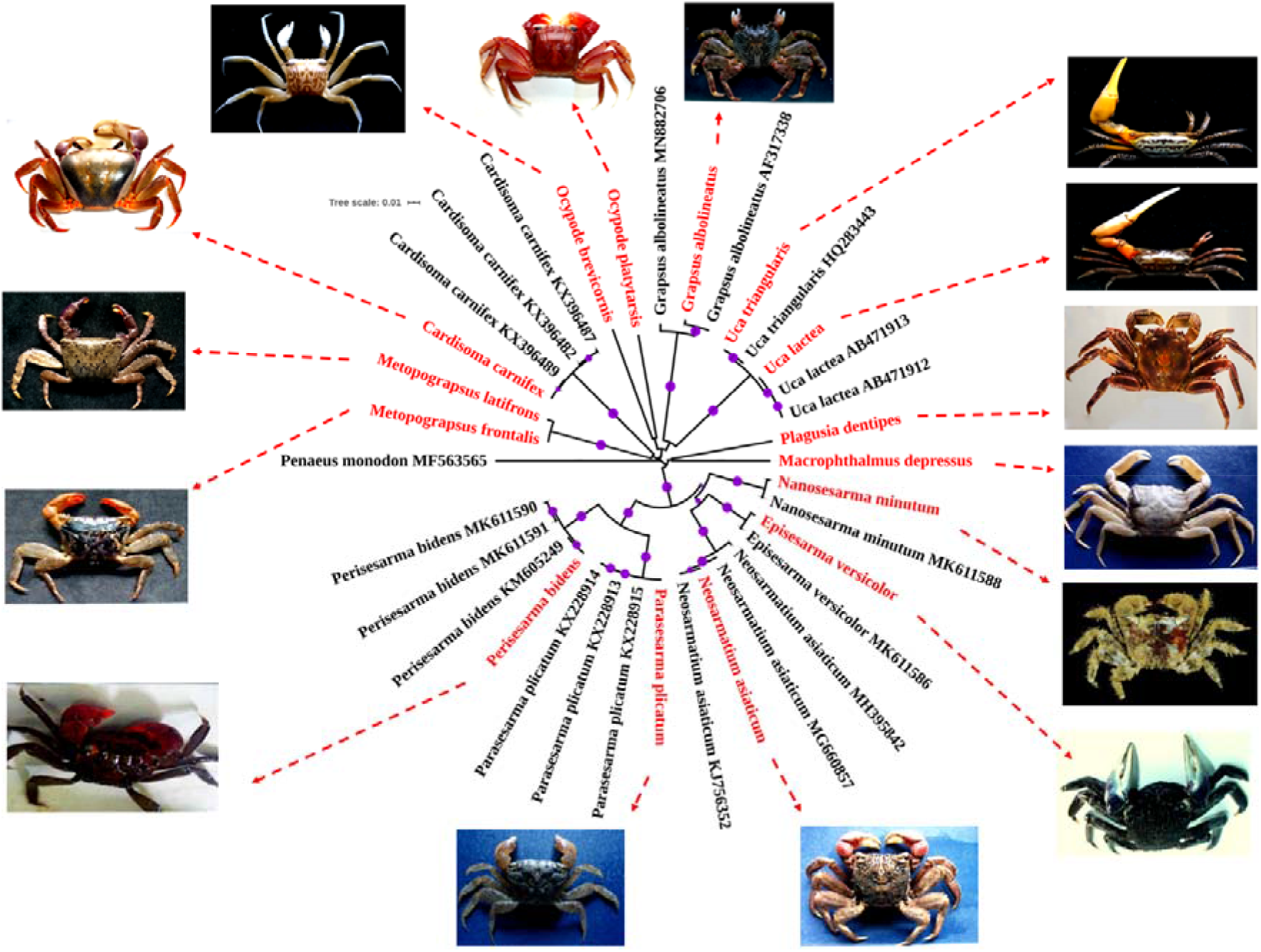
Neighbour-joining tree was drawn using Kimura-2 parametric distance model. Species marked in red and black were sequences of this study and reference Genbank sequences (with accession numbers), respectively. Bootstrap values >70 were shown in purple circles. *Penaeus monodon* (Genbank Acc. No.: 563565) used as an out-group. Species without reference sequences in the phylogram indicates the first-time barcoded species.

The SPM clade grouped all Sesarmidae members *(Perisesarma bidens, Parasesarma plicatum, Neosarmatium asiaticum, Episesarma versicolor* and *Nanosesarma minutum*) into one major clade, and members of Plagusiidae (*Plagusia dentipes*), Macrophthalmidae (*Macrophthalmus depressus*) in the neighbouring clade. GG clade also separated Gecarcinidae (*Cardisoma carnifex*) from other members of the Grapsidae (*Metopograpsus frontalis, M. latifrons*). However, OG clade grouped Ocypodidae members (*Uca lactea, U. triangularis, Ocypode brevicornis* and *O. platytarsis*) into single clade while placing a Grapsidae member (*Grapsus albolineatus*) in the neighbouring clade. The reason *Grapus albolineatus* does not clade group with other members of the same family (in GG clade) is unknown, and therefore the phylogenetic relationships between Grapsidae members which need further investigation.

The K2P distance showed that *Grapsus albolineatus* had been genetically closer to *Uca* spp. (0.21) excluding the *Ocypode* spp. (0.22) between members of the OG clade (table S4). When K2P distance was compared between *G. albolineatus* and other Grapsidae members *(Metopograpsus latifrons* and *M. frontalis*), the values were slim (~0.2) (**Table S4**). The mis-placement of *G. albolineatus* in NJ tree and higher genetic variations among the members of Grapsidae may deserve further investigation. The average K2P distances among all the brachyuran crab species barcoded in this study were 0.2.

## 4. Discussion

Diversity is a measure of the complexity of the community structure (example: Cisneros et al., 2011; Menta, 2012) and the mangroves ecosystem’s biodiversity changes due to physical, chemical and biological factors (Kathiresan and Bingham, 2001; Carugati et al., 2018). High diversity within mangrove ecosystems suggest that a community is balanced, stable and responsive (Carugati et al., 2018). Many habitats are highly dependent on their faunal communities which are dominated by brachyuran crabs. Brachyuran crabs are among the most important components of both natural and artificially created mangrove ecosystems (Cannicci et al., 2008, Kristensen et al., 2008), but their occurrences and preferences in mangrove ecosystems artificially created are not well established.

Unlike Tampa Bay, Florida, where tidal salt marshes created that were naturally transposed into mangrove wetlands (Osland et al., 2012), Vellar estuary mangroves were planned and resurrected from mangrove saplings (see, Ajmal Khan et al., 2005). Vellar estuary mangroves played an important role in Tsunami mitigation during 2004 (Kathiresan & Rajendran, 2005). During flooding and competent larvae settling, brachyuran crab larvae are transported into mangrove ecosystems (Epifanio, 1995). Most brachyuran crab larvae were highly competent in locating conspecific adults colonizing the forest area (Cannicci et al., 2019). *Parasesarma capensis* is more abundant in the *Rhizophora mucronata* characterized areas (Fratini et al., 2019). During settlement and colonization, different species of brachyuran crab larvae prefers different mangrove species (Cannicci et al., 2019).

### 4.1. Mangroves associated brachyuran crab diversity

It is not a comprehensible inference that abundance would contribute to the number of species per unit area. This relation depends on the community level of equity (Glover et al., 2001). In the present study, a total of 2844 crabs representing 35 species representing 8 families (*viz*., Sesarmidae, Portunidae, Ocypodidae, Grapsidae, Dotillidae, Varunidae, Macrophthalmidae and Gecarcinidae) were collected, where over 70% of total abundance was dominated by 4 species of crab, i.e., *Uca lactae, U. triangularis, Selatium brockii*, and *Neosarmatium asiaticum*.

Station 3M had maximum species diversity that is likely due to increased physical and environmental stability, as stated by Levin and Gage (1998), Ganesh, (2003), Ajmal khan et al. (2005), Bijukumar et al. (2007) and Praveenkumar *et al*. (2013). Though it was unclear how a mangrove species in a mixed mangrove community influences the colonization and settlement of a crab species, such trends were also observed in Brazilian mangroves (Colpo et al., 2011). However, this does not alter the observed fact that the type of mangrove species (or community) affects the structure and composition of the brachyuran crab community, since the pattern of distribution of crabs was due to the presence or absence of a particular mangrove species observed in this study.

Species such as *Uca lactea, U. triangularis* and *Neosarmatium asiaticum* have been found to be consistently abundant at all of the study area stations. The bar plot and the shade plot thus clearly explained the species-richness of brachyuran crabs of Vellar mangrove, in agreement with earlier works by Bijukumar et al., (2007) and Bharadhirajan (2012). *Selatium brockii* abundances have been substantially reduced at non-mangrove stations (7N and 2N), exposing this species’ heavy reliance on mangrove vegetation.

### 4.2. Comparing brachyuran crab abundances in artificial mangroves between 2005 and 2015

It took about 15 years for 8 species of brachyuran crabs to colonize and the remaining species were colonized in the next 10 years, when compared with the near-by Pitchavaram ecosystem’s brachyuran crab species composition. This colonization pattern shows that the effects of transport and settlement are cumulative over time. In the present analysis the sampled stations VI and VII in Ajmal Khan et al, (2005) were 3M and 4A. The present study found that, during 2005, the abundance of 8 species (*Selatium brockii, Episesarma mederi, Nanosesarma minutum, N. batavicum, Grapsus tenuicrustatus, Uca lactea, U. triangularis*, and *Dotilla myctiroides*) colonizing 3M and 4A almost doubled (with few exceptions, such as *G. tenuicrustatus* missing from 3M). The H diversity index also increased by more than 4 times in 3M and 4A, when compared to 2005 sampling (Ajmal Khan et al., 2005). That is, the H diversity index was 8 and 5 during 2005 for the stations 3M and 4A, respectively which is 33 and 22, respectively in the present analysis. This may be clarified due to variations in the mangrove composition observed over various time period at the given sampling station. That is, during 2005, station 3M consisted of *Rhizopora apiculata, Avicennia marina, Acanthus illicifolius* and station 4A consisting of *Avicennia marina* and *Acanthus illicifolis* (Ajmal Khan et al., 2005). Present study found that, station 3M is composed of *Avicennia marina, A. officinalis, Rhizophora apiculata, R. mucronata* and *R. annamalayana*. The station 4A only composed with *Avicennia marina*. Herbaceous macrophytes may have played a role in influencing the mangrove composition by trapping mangrove propagules from local supply at the given station (McKee et al., 2007; Peterson and Bell, 2012; Lewis, 2005). Therefore, one should be aware of changes in the composition of mangrove spatially and temporally (in the artificially created mangrove patches) in order to identify changes in the composition of the inhabitation brachyuran crab community over time.

### 4.3. Why DNA barcoding selective brachyuran crab species?

While 8 crab species occurred during 2005 were precisely identified (Ajmal Khan et al., 2005), taxonomic ambiguity was faced in this study (example, in Gecarcinidae; *Cardisoma carnifex)*. The key explanation for this was the non-fusion of the supra- and infra-orbital margins along the outer edges which can be called incomplete orbits (Ng et al., 2008). The challenging problem was also the distinctive features such as the inclusion versus exclusion of the antennae from the orbital hiatus which is necessary to differentiate between *Pachygrapsus* (includes antennae) from *Metopograpsus* (excludes antennae) (Poupin et al. 2005). Including among the members of Ocypodidae, while the number of features can be used to classify species within this genus Ocypode, many of these are not generally useful for establishing natural groupings (Sakai and Türkay, 2013) and the uncertainty and dispute over the proposed generic, subgeneric, and specific taxonomy of the genus *Uca* (Rosenberg, 2001) has been witnessed.

It was tough to differentiate the members of Plagusiidae and Grapsidae. While the crabs, *Plagusia dentipes* (Plagusiidae) and *Grapsus albolineatus* (Grapsidae) belong to the two separate families, earlier they belonged to the Plagusinae and Grapsinae subfamilies under Grapsidae family, respectively. The subfamilies have now been raised to family-level. Consequently, difficulties in identifying those species using morphological characters are understandable, and molecular identification is necessary. Sesarmidae- the keystone species of mangrove communities (Smith et al., 1991; Lee, 1998), even after many taxonomic revisions (e.g., Davie, 1992; 1994; Schubart et al., 2009; Davie, 2012), there are still several ambiguities and taxonomic issues that call for further studies (see notes on Sesarmidae, Ng et al., 2008). Even after splitting into 5 sub-genera (Barnes, 2010), taxonomic problems prevailed in the genus *Macrophthalmus* was solved (Ng et al., 2008, Davie, 2009).

### 4.4. DNA barcoding and pairwise analysis

BLAST analysis revealed the species; *Plagusia dentipes, Ocypode platytarsis, O. brevicornis, Metopograpsus frontalis*, and *M. latifrons* and the genus *Macrophthalmus* sp. *(M. depressus)* were barcoded for the first time. We realised that as many more commonly available brachyuran crab species lacked reference barcodes in Ganbank, DNA barcoding needs veen stronger push globally and locally. For example, while *Macrophthalmus depressus* was described as early as the 1830’s, (Rüppell, 1830), Genbank was absent from the barcode sequence of this genera as a whole.

Minimal genetic differentiation is expected within the SPM clade between the *Periseserma bidens* and *Parasesarma plicatum*, as most species of *Parasesarma* (De Man, 1895) was transferred from *Perisesarma* (De Man, 1895) (see Shahdadi and Schubart 2017). The distinction of *Parasesarma* and *Perisesarma* was previously due to the presence or absence of a distinct epibranchial tooth, and while this was long suspected of being artificial in nature, Shahdadi and Schubart (2015, 2017) finally demonstrated that character was phylogenetically uninformative. Like most sesarmid crabs, the *Parasesarma* genus has marine planktonic larvae (see Flores et al., 2002; Guerao et al., 2004; Lago 1993) and thus a potential for high dispersal capability. Recognition of distinct brachyuran crab family Macrophthalmidae (Dana, 1851) from Ocypodidae (Rafinesque, 1815) on the basis of morphological and molecular evidence (see Kitaura et al., 2002; Mendoza & Ng, 2007; Ng et al., 2008) was reaffirmed in this analysis as *Macrophthalmus depressus* was grouped within the SPM clade, not in OG clade where Ocypodidae members were grouped together. The Grapsidae (MacLeay, 1838) crab family, was originally a large group of thoracotreme crabs, with over 400 species in over 50 genera within the superfamily Grapsoidea (see Guinot, 1978; Bowman & Abele, 1982).

GG clade seperated Gecarcinidae (*Cardisoma carnifex*) from other members of Grapsidae (*Metopograpsus frontalis, M. latifrons*), excerpted from *Grapsus albolineatus* grouped within the OG clade. Grouping of Gecarcinidae and Grapsidae was expected as they were under single superfamily Grapsoidea which was also supported previously using molecular data (Schubart et al., 2000; 2002). Grapsidae phylogeny may require reinvestigation as OG clade contained the members of Ocypodidae (*Uca lactea, U. triangularis, Ocypode brevicornis* and *O. platytarsis*), while the member of Grapsidae (*Grapsus albolineatus*) was distanced in the neighbouring clade.

Compared with the Ocypodoidea the Grapsoidea does not seem to be reciprocally monophyletic (see Schubart, Neigel & Felder, 2000b; Kitaura, Wada & Nishida, 2002; Schubart et□al., 2006). As such, Schubart et□al. (2006) proposed that it would be better to avoid using these taxa until monophyletic superfamilies could be re-defined and in the meantime simply refer to the entire group as Thoracotremata. The genus *Grapsus* spp. was paraphyletic and would not necessarily be accommodated in the Grapsidae family clade (Ip et al., 2015). The higher similarity between *Metograpsus frontalis* and *M. latifrons* was observed in the previous analysis consisting of 5 different *Metapograpsus* spp. (Ip et al., 2015). The multi-barcode markers approach also verified that *Grapsus albolineatus* would not be identified with members of Grapsidae (see IP et al., 2015). The multiple barcode phylogeny also revealed that *G. albolineatus* was genetically similar to *Pachygrapsus fakaravensis*. Therefore this unique species (*G. albolineatus*) can need further phylogenetic re-examination.

Universal barcode gap proposal would be mandatory to delineate brachyuran crabs from meta- and environmental barcoding outcomes (Weigand et al., 2019). The average genetic distances (K2P) for all brachyuran crab species in this study were 0.2. Proposing a 2% gap cannot be feasible (based on overall pairwise distance data from this present study) as it was recognised that some species of Plagusiidae (*Perisesarma guttatum*) contain 2.75% K2P distance within various haplotypes along the East African coast (Silva et al., 2010). As new species *Parasesarma austrawati* (in northern Australian mangroves (Shahdadi et al., 2019)) had 4% genetic distances among other species of *Paraseserma* spp., the genetic distance values of 3% species barcode gap, previously proposed by Hebert et al., (2003), may not work for brachyuran crabs. For brachyuran crab species delineation, family specific COI barcode gaps should be defined.

## 5. Conclusion

A combination of morphological and molecular data will certainly give us the most information (see Klaus et al., 2009) and would allow us to make a major leap towards our ultimate goal of reconstructing the decapod Tree of Life. First time barcodes produced in this study shows the needs to improve the barcode species coverage in reference libraries. It should also be borne in mind that DNA barcoding reimbursement is not restricted to taxonomic or systematic research only. The emergence of modern high-throughput sequencing technologies will in the near future significantly change applications and surveys for bio-monitoring (Fonseca et al., 2010; Hajibabaei et al., 2011; Leray et al., 2015). This will make reference datasets such as ours important for the identification of barchyuran crabs. The current research found it would take approximately 25 years for the Vellar mangroves to absorb almost all brachyuran crab species of Pichavaram mangroves. Using the array of diversity and species estimator indices resulted in useful output data for the period needed for colonization, and preferences of artificially developed mangrove by brachyuran crabs. This is a bench mark data for marine policy makers, coastal ecosystem designers and climate researchers.

## 6. Acknowledgement

First author (GM) thank Ministry of Earth Sciences (Centre for Marine Living Resources and Ecology) for the financial grand (No. 10-IT IS/17/2012 dt.01/10/2012). The second author (CP) thank the Department of Science and Technology’s INSPIRE Fellowship (IF10431), India and China Postdoctoral Research Foundation’s National Postdoctoral fellowship (0050-K83008), China for the partial financial assistance. Authors are grateful to the anonymous reviewer’s for improving the quality of the manuscript.

## 7. Declarations

Funding (acknowledged in the acknowledment section)

Conflicts of interest/Competing interests: Authors declares no Conflicts/Competing interests.

Ethics approval: Not applicable

Consent to participate: Not applicable

Consent for publication: Not applicable

Availability of data and material: Supplementary materials included

Code availability: Not applicable

### Authors’ contributions

GM and CP were involved in designing, field collection and data acquisition and interpretation, CP, JV, SRP and NP was involved in fund acquisition, data interpretation and draft preparation.

## References

Ajmal Khan, S., Raffi, S. M., Lyla, P, S. (2005). Brachyuran crab diversity in natural (Pitchavaram) and artificially developed mangroves (Vellar estuary). Current Science. 88(8): 1316–1324

Alongi, D. M. (2002). Present state and future of the world’s mangrove forests. Environmental Conservation. 29, 331–349. doi:10.1017/s0376892902000231.

Alongi, D. M. (2012). Carbon sequestration in mangrove forests. Carbon Management. 3, 313–322. doi:10.4155/cmt.12.20.

Alongi, D. M. (2015). The Impact of Climate Change on Mangrove Forests. Current Climate Change Reports. 1, 30–39. doi:10.1007/s40641-015-0002-x.

Altschul, S. (1997). Gapped BLAST and PSI-BLAST: a new generation of protein database search programs. Nucleic Acids Research. 25, 3389–3402. doi:10.1093/nar/25.17.3389.

Carugati, L., Gatto, B., Rastelli, E., Lo Martire, M., Coral, C., Greco, S., et al. (2018). Impact of mangrove forests degradation on biodiversity and ecosystem functioning. Scientific Reports. 8. doi:10.1038/s41598-018-31683-0.

Ballard JWO, Rand DM. The population biology of mitochondrial DNA and its phylogenetic implications. Annu Rev Ecol Evol S 2005; 36: 621–642.

Ballard JWO, Whitlock MC. The incomplete natural history of mitochondria. Mol Ecol 2004; 13:729–744.

Beng, K.C., Corlett, R.T. Applications of environmental DNA (eDNA) in ecology and conservation: opportunities, challenges and prospects. Biodivers Conserv 29, 2089–2121 (2020). https://doi.org/10.1007/s10531-020-01980-0

Bernt M, Braband A, Schierwater B, Stadler PF. Genetic aspects of mitochondrial genome evolution. Mol Phylogenet Evol 2013; 69: 328–338.

Biella, P., & Galimberti, A. (2020). The spread of <em>Bombus haematurus</em> in Italy and its first DNA barcode reference sequence. Fragmenta Entomologica, 52(1), 67–70. https://doi.org/10.4081/fe.2020.413

Bowmann T.E., and Abele, L.G. (1982). Classification of the recent Crustacea. in The biology of Crustacea 1: systematics, the fossil record and biogeography, ed. L.G. Abele (New York, Academic Press), 1–27.

Brandão, M. C., Freire, A. S., & Burton, R. S. (2016). Estimating diversity of crabs (Decapoda: Brachyura) in a no-take marine protected area of the SW Atlantic coast through DNA barcoding of larvae. Systematics and Biodiversity, 14(3), 288–302. https://doi.org/10.1080/14772000.2016.1140245

Brown, G. R. (2019). First DNA barcode for the enigmatic Leiobunum sp. A (Opiliones). Arachnology, 18(2), 94. https://doi.org/10.13156/arac.2018.18.2.94

Bucklin A, Steinke D, Blanco-Bercial L. DNA barcoding of marine Metazoa. Ann Rev Mar Sci 2011; 3:471–508.

Cannicci, S., Fusi, M., Cimó, F., Dahdouh-Guebas, F., and Fratini, S. (2018). Interference competition as a key determinant for spatial distribution of mangrove crabs. BMC Ecology. 18. doi:10.1186/s12898-018-0164-1.

Cannicci, S., Mostert, B., Fratini, S., McQuaid, C., and Porri, F. (2019). Recruitment limitation and competent settlement of sesarmid crab larvae within East African mangrove forests. Marine Ecology Progress Series. 626, 123–133. doi:10.3354/meps13062.

Chen Y-Y, Lin H-C, Chan BKK. Description of a new species of coral-inhabiting barnacle, Darwiniella angularis sp. n. (Cirripedia, Pyrgomatidae) from Taiwan. ZooKeys 2012; 214: 43–74.

Colpo, K.D., Chacur, M.M., Guimarães, F.J. et al. Subtropical Brazilian mangroves as a refuge of crab (Decapoda: Brachyura) diversity. Biodivers Conserv 20, 3239–3250 (2011). https://doi.org/10.1007/s10531-011-0125-x

De Grave S, Chu KH, Chan T-Y. On the systematic position of Galatheacaris abyssalis (Decapoda: Galatheacaridoidea). J Crustacean Biol 2013; 30: 521–527.

Dev Roy, M.K. (2008). An annotated checklist of mangrove and coral reef inhabiting Brachyuran crabs of India. Rec. Zool. Surv. India, Occ. Paper. 289:1–212. (Published by the Director, Zool. Surv. India, Kolkata)

Egli, D., Olckers, T., Willows-Munro, S., & Harvey, K. (2020). DNA barcoding of endophagous immature stages elucidates the host-plant affinities of insects associated with the invasive Senecio madagascariensis in its native range in South Africa. Biological Control, 145, 104245. https://doi.org/10.1016/j.biocontrol.2020.104245

Ellison, J., and Zouh, I. (2012). Vulnerability to Climate Change of Mangroves: Assessment from Cameroon, Central Africa. Biology. 1, 617–638. doi:10.3390/biology1030617.

Elwin, A., Bukoski, J.J., Jintana, V. et al. Preservation and recovery of mangrove ecosystem carbon stocks in abandoned shrimp ponds. Sci Rep 9, 18275 (2019). https://doi.org/10.1038/s41598-019-54893-6

Epifanio, C.E. (1995). Transport of blue crab (Callinectes sapidus) larvae in the waters off mid Atlantic states. Bulletine of Marine Science. 57, 713–725.

FAO, (2007). Food and Agriculture Organization of the United Nations In The world’s mangroves 1980–2005. Forest Resources Division, FAO, Rome: FAO Forestry Paper 153.

Feroz K S., Sanker G., Prasannakumar C (2014) Linking eggs and adults of Argulus spp. using mitochondrial DNA barcodes. Mitochondrial DNA 10: 1–5. https://doi.org/10.3109/19401736.2014.987269

Flores, A. A. V., Paula, J., and Saraiva, J. (2002). Sexual Maturity, Reproductive Cycles, and Juvenile Recruitment of Perisesarma Guttatum (Brachyura, Sesarmidae) at Ponta Rasa Mangrove Swamp, Inhaca Island, Mozambique. Journal of Crustacean Biology. 22, 143–156. doi:10.1163/20021975-99990217.

Folmer, O., Black, M., Hoeh, W., Lutz, R., and Vrijenhoek, R. (1994). DNA primers for amplification of mitochondrial cytochrome c oxidase subunit I from diverse metazoan invertebrates. Molecular Marine Biology and Biotechnology. 3, 294–299.

Fratini, S., Cannicci, S., Porri, F., Innocenti, G. (2019). Revision of the Parasesarma guttatum species complex reveals a new pseudo-cryptic species in South East African mangroves. Invertebrate Systematics. 33, 208–224.

Fratini, S., Vigiani, V., Vannini, M., and Cannicci, S. (2004). Terebralia palustris (Gastropoda; Potamididae) in a Kenyan mangal: size structure, distribution and impact on the consumption of leaf litter. Marine Biology. 144, 1173–1182. doi:10.1007/s00227-003-1282-6.

Gilman, E. L., Ellison, J., Duke, N. C., and Field, C. (2008). Threats to mangroves from climate change and adaptation options: A review. Aquatic Botany. 89, 237–250. doi:10.1016/j.aquabot.2007.12.009.

Giri, C., Ochieng, E., Tieszen, L. L., Zhu, Z., Singh, A., Loveland, T., et al. (2010). Status and distribution of mangrove forests of the world using earth observation satellite data. Global Ecology and Biogeography. 20, 154–159. doi:10.1111/j.1466-8238.2010.00584.x.

Guerao, G. (2004). Complete larval and early juvenile development of the mangrove crab Perisesarma fasciatum (Crustacea: Brachyura: Sesarmidae) from Singapore, with a larval comparison of Parasesarma and Perisesarma. Journal of Plankton Research. 26, 1389–1408. doi:10.1093/plankt/fbh127.

Guinot, D. (1978). Principes d’une classification évolutive des Crustacés Décapodes Brachyoures. Bulletin Biologique de la France et de la Belgique. 112, 211–292.

Gunalan B and Prasannakumar C (2014) Report of Gnathophausia ingens (Dohrn, 1870) from bathypelagic region of Bay of Bengal, corroborated by DNA barcoding and 18S rRNA gene sequencing. Indian J. Fish., 61(4): 123–126.

Hebert PDN, Cywinska A, Ball SL, deWaard JR. Biological identifications through DNA barcodes. Proc Biol Sci 2003b; 270: 313–321.

Hebert PDN, Ratnasingham S, deWaard JR. Barcoding animal life: cytochrome c oxidase subunit 1 divergences among closely related species. Proc Biol Sci 2003a; 270: S96–S99.

Hebert, P. D. N., Cywinska, A., Ball, S. L., and deWaard, J. R. (2003). Biological identifications through DNA barcodes. Proceedings of the Royal Society of London. Series B: Biological Sciences. 270, 313–321. doi:10.1098/rspb.2002.2218.

Hemalatha, A., Mohammed Esa, S. A. R., Suresh, M., Thajuddin, N., & Anantharaman, P. (2016). Identification of Odontella aurita by rbcL gene sequence – a high antibacterial potential centric marine diatom. Mitochondrial DNA Part A, 28(5), 655–661. https://doi.org/10.3109/24701394.2016.1166222

Hsiang, L.L. Mangrove conservation in Singapore: A physical or a psychological impossibility?. Biodiversity and Conservation 9, 309–332 (2000). https://doi.org/10.1023/A:1008993417327

Igulu, M. M., Nagelkerken, I., Dorenbosch, M., Grol, M. G. G., Harborne, A. R., Kimirei, I. A., et al. (2014). Mangrove Habitat Use by Juvenile Reef Fish: Meta-Analysis Reveals that Tidal Regime Matters More than Biogeographic Region. PLoS ONE. 9, e114715. doi:10.1371/journal.pone.0114715.

Ip, B. H. Y., Schubart, C. D., Tsang, L. M., and Chu, K. H. (2015). Phylogeny of the shore crab family Grapsidae (Decapoda: Brachyura: Thoracotremata) based on a multilocus approach. Zoological Journal of the Linnean Society. 174, 217–227. doi:10.1111/zoj.12235.

Kathiresan, K., and Bingham, B. L. (2001). ‘Biology of mangroves and mangrove Ecosystems’, in Advances in Marine Biology. 81–251. doi:10.1016/s0065-2881(01)40003-4.

Kathiresan, K., and Rajendran, N. (2005). Coastal mangrove forests mitigated tsunami. Estuarine, Coastal and Shelf Science. 65, 601–606. doi:10.1016/j.ecss.2005.06.022.

Kelnarova, I., Jendek, E., Grebennikov, V. V., & Bocak, L. (2018). First molecular phylogeny of Agrilus (Coleoptera: Buprestidae), the largest genus on Earth, with DNA barcode database for forestry pest diagnostics. Bulletin of Entomological Research, 109(2), 200–211. https://doi.org/10.1017/s0007485318000330

Khalaji-Pirbalouty V, Raupach MJ. A new species of Cymodoce Leach, 1814 (Crustacea: Isopoda: Sphaeromatidae) based on morphological and molecular data, with a key to the Northern Indian Ocean species. Zootaxa 2014; 3826: 230–254.

Khan, S. A., Lyla, P. S., John, B. A., Kuamr, C. P., Murugan, S., & Jalal, K. C. A. (2010). DNA Barcoding of Stolephorus indicus, Stolephorus commersonnii and Terapon jarbua of Parangipettai Coastal Waters. Biotechnology (Faisalabad), 9(3), 373–377. https://doi.org/10.3923/biotech.2010.373.377

Khan, S. A., Prasannakumar C, Lyla P. S., Murugan S (2011) Identifying Marine fin fishes using DNA barcodes. Current Science, vol. 101 (9): 1152–1154.

Kimura M. (1980). A simple method for estimating evolutionary rate of base substitutions through comparative studies of nucleotide sequences. Journal of Molecular Evolution 16:111–120.

Kitaura, J., Wada, K., and Nishida, M. (2002). Molecular Phylogeny of Grapsoid and Ocypodoid Crabs with Special Reference to the Genera Metaplax and Macrophthalmus. Journal of Crustacean Biology. 22, 682–693. doi:10.1163/20021975-99990281.

Krishnamurthy, K. P., Francis, R.A. A critical review on the utility of DNA barcoding in biodiversity conservation. Biodivers Conserv 21, 1901–1919 (2012). https://doi.org/10.1007/s10531-012-0306-2

Kristensen, E., Bouillon, S., Dittmar, T., and Marchand, C. (2008). Organic carbon dynamics in mangrove ecosystems: A review. Aquatic Botany. 89, 201–219. doi:10.1016/j.aquabot.2007.12.005.

Lago, R. P. (1993). Larval Development of Sesarma guttatum A. Milne Edwards (Decapoda: Brachyura: Grapsidae) Reared in the Laboratory, with Comments on Larval Generic and Familial Characters. Journal of Crustacean Biology. 13, 745. doi:10.2307/1549105.

Lee, S. Y., Primavera, J. H., Dahdouh-Guebas, F., McKee, K., Bosire, J. O., Cannicci, S., et al. (2014). Ecological role and services of tropical mangrove ecosystems: a reassessment. Global Ecology and Biogeography. 23, 726–743. doi:10.1111/geb.12155.

Lewis, R. R., III (2005). Ecological engineering for successful management and restoration of mangrove forests. Ecological Engineering. 24, 403–418. doi:10.1016/j.ecoleng.2004.10.003.

Li, J., Cui, Y., Jiang, J. et al. Applying DNA barcoding to conservation practice: a case study of endangered birds and large mammals in China. Biodivers Conserv 26, 653–668 (2017). https://doi.org/10.1007/s10531-016-1263-y

Lovelock, C. E., Cahoon, D. R., Friess, D. A., Guntenspergen, G. R., Krauss, K. W., Reef, R., et al. (2015). The vulnerability of Indo-Pacific mangrove forests to sea-level rise. Nature. 526, 559–563. doi:10.1038/nature15538.

Madhavan, A., Silvester, R., Prabhakaran, M. P., Naderloo, R., Radhakrishnan, C. K., & Menon, N. R. (2020). First barcode of Ryphila cancellus (Herbst, 1783), from the southwest coast of India. Regional Studies in Marine Science, 33, 100910. https://doi.org/10.1016/j.rsma.2019.100910

Manikantan, G., PrasannaKumar, C., Vijaylaxmi, J., Pugazhvendan, S. R., & Prasanthi, N. (2020). Diversity, phylogeny, and DNA barcoding of brachyuran crabs in artificially created mangrove environments. Cold Spring Harbor Laboratory. https://doi.org/10.1101/2020.09.07.286823

Markert A, Raupach MJ, Segelken-Voigt A, Wehrmann A. Molecular identification and morphological characteristics of native and invasive Asian brush-clawed crabs (Crustacea: Brachyura) from Japanese and German coasts: Hemigrapsus penicillatus (De Haan, 1835) versus Hemigrapsus takanoi Asakura & Watanabe 2005. Org Divers Evol 2014; 14: 369–382.

McKee, K. L., Rooth, J. E., and Feller, I. C. (2007). Mangrove recruitment after forest disturbance is facilitated by herbaceous species in the Caribbean. Ecological Applications. 17, 1678–1693. doi:10.1890/06-1614.1.

McLeod, E. and Salm, R.V. (2006) Managing Mangroves for Resilience to Climate Change. The International Union for the Conservation of Nature and Natural Resources (IUCN), Gland, Switzerland.

Meiklejohn, K. A., Wallman, J. F., & Dowton, M. (2013). DNA Barcoding Identifies all Immature Life Stages of a Forensically Important Flesh Fly (Diptera: Sarcophagidae). Journal of Forensic Sciences, 58(1), 184–187. https://doi.org/10.1111/j.1556-4029.2012.02220.

Mendoza, J. C. E. and Ng, P. K. L. (2007). Macrophthalmus (Euplax) H. Milne-Edwards, 1852, a valid subgenus of ocypodoid crab (Decapoda: Brachyura: Macrophthalmidae), with description of a new species from the Philippines. Journal of Crustacean Biology. 27(4), 860–870.

Menta, C. (2012). ‘Soil Fauna Diversity - Function, Soil Degradation, Biological Indices, Soil Restoration’, Biodiversity Conservation and Utilization in a Diverse World, ed. Dr. Gbolagade Akeem Lameed (InTech). doi:10.5772/51091.

Ng, P. K. L., Guinot, D. and Davie, P. J. F. (2008). Systema Brachyurorum: Part I. An annotated checklist of the extant brachyuran crabs of the world. The Raffles Bulletin of Zoology. 17, 1–286.

Ortega Cisneros, K., Smit, A. J., Laudien, J., and Schoeman, D. S. (2011). Complex, Dynamic Combination of Physical, Chemical and Nutritional Variables Controls Spatio-Temporal Variation of Sandy Beach Community Structure. PLoS ONE 6, e23724. doi:10.1371/journal.pone.0023724.

Osland, M. J., Spivak, A. C., Nestlerode, J. A., Lessmann, J. M., Almario, A. E., Heitmuller, P. T., et al. (2012). Ecosystem Development After Mangrove Wetland Creation: Plant–Soil Change Across a 20-Year Chronosequence. Ecosystems. 15, 848–866. doi:10.1007/s10021-012-9551-1.

Palanisamy, S. K., PrasannaKumar, C.*, Paramasivam, P., & Sundaresan, U. (2020). DNA barcoding of horn snail Telescopium telescopium (Linnaeus C, 1758) using mt-COI gene sequences. Regional Studies in Marine Science, 35, 101109. https://doi.org/10.1016/j.rsma.2020.101109

Peterson, J. M., and Bell, S. S. (2012). Tidal events and salt-marsh structure influence black mangrove (Avicennia germinans) recruitment across an ecotone. Ecology. 93, 1648–1658. doi:10.1890/11-1430.1.

Polidoro, B. A., Carpenter, K. E., Collins, L., Duke, N. C., Ellison, A. M., Ellison, J. C., et al. (2010). The Loss of Species: Mangrove Extinction Risk and Geographic Areas of Global Concern. PLoS ONE. 5, e10095. doi:10.1371/journal.pone.0010095.

PrasannaKumar C, Akbar John B, Ajmal Khan S, Lyla P S, and Jalal K C A (2012) Limit of DNA Barcode in Delineating Penaeus Monodon and in its Developing Stages. Sains Malaysiana 41(12): 1527–1533.

Prasannakumar, C., Iyyapparajanarasimapallavan, G., Rahman M. A. U., Mohanchander, P., Sudhakar, T. (2020b) Variability in the diet diversity of catfish highlighted through DNA barcoding. Cold Spring Harbor Laboratory. https://doi.org/10.1101/2020.09.18.268888

Prasannakumar, C., Manikantan, G., Vijaylaxmi, J., Gunalan, B., Manokaran, S., & Pugazhvendan, S. R. (2020c). Strengthening marine amphipod DNA barcode libraries for environmental monitoring. Cold Spring Harbor Laboratory. https://doi.org/10.1101/2020.08.26.268896

PrasannaKumar, C., Rethinavelu, S., & Sadaiappan, B. (2020a). First barcodes of Bathynomus kensleyi (Lowry & Dempsey, 2006) and Bathynomus decemspinosus (Shih, 1972) from the Southeast coast of India. Regional Studies in Marine Science, 40, 101489. https://doi.org/10.1016/j.rsma.2020.101489

Prasanthi, N., Prasannakumar, C., Annadurai, D., & Mahendran, S. (2020). Identifying seaweeds species of Chlorophyta, Ochrophyta and Rhodophyta using DNA barcodes. Cold Spring Harbor Laboratory. https://doi.org/10.1101/2020.08.30.274456

Rahman A M, Khan S A, Lyla P S, PrasannaKumar C (2013) DNA Barcoding Resolves Taxonomic Ambiguity in Mugilidae of Parangipettai Waters (Southeast Coast of India). Turkish Journal of Fisheries and Aquatic Sciences 13: 321–330.

Rahman, M. A. U., PrasannaKumar, C., Mohanchander, P., Manikantan, G., Ajmal Khan, S. and Lyla, P.S. (2019) Identification of eggs, larva and adults of Scylla serrata (Forsskal, 1775) using DNA barcodes. Journal of Aquatic Biology & Fisheries. 7: 24–30.

Riehl T, Kaiser S. Conquered from the deep sea? A new deep-sea isopod species from the Antarctic shelf shows pattern of recent colonization. PLoS One 2012; 7: e49354.

Rüppell, E. S. (1830). Beschreibung und Abbildung von 24 Arten kurzschwänzigen Krabben, als Beitrag zur Naturgeschichte des rothen Meeres. H.L. Brönner, Frankfurt. 1–28, 1–6.

Saenger, P. (2002). In Mangrove ecology, Silviculture and conservation. Dordrecht: Kluwer Academic Publishers.

Sahu, S. K., Singh, R., & Kathiresan, K. (2016). Multi-gene phylogenetic analysis reveals the multiple origin and evolution of mangrove physiological traits through exaptation. Estuarine, Coastal and Shelf Science, 183, 41–51. https://doi.org/10.1016/j.ecss.2016.10.021

Salmo, S. G., III, Lovelock, C., and Duke, N. C. (2013). Vegetation and soil characteristics as indicators of restoration trajectories in restored mangroves. Hydrobiologia. 720, 1–18. doi:10.1007/s10750-013-1617-3.

Sandilyan, S., Kathiresan, K. Mangrove conservation: a global perspective. Biodivers Conserv 21, 3523–3542 (2012). https://doi.org/10.1007/s10531-012-0388-x

Sathiadhas, R., George, J. P., Jayasurya, P. K., Mathew, Ansy. (2005). Economic Importance of Mangroves, Afforestation and Reclamation. Mangrove ecosystems: A manual for the assessment of biodiversity. (CMFRI Special Publication), 83, 215–218. http://eprints.cmfri.org.in/id/eprint/4050.

Schubart, C. D., Cannicci, S., Vannini, M., and Fratini, S. (2006). Molecular phylogeny of grapsoid crabs (Decapoda, Brachyura) and allies based on two mitochondrial genes and a proposal for refraining from current superfamily classification. Journal of Zoological Systematics and Evolutionary Research. 44, 193–199. doi:10.1111/j.1439-0469.2006.00354.x.

Schubart, C. D., Cuesta, J. A., and Felder, D. L. (2002). Glyptograpsidae, a New Brachyuran Family from Central America: Larval and Adult Morphology, and a Molecular Phylogeny of the Grapsoidea. Journal of Crustacean Biology. 22, 28–44. doi:10.1163/20021975-99990206.

Schubart, C. D., Cuesta, J. A., Diesel, R., and Felder, D. L. (2000a). Molecular Phylogeny, Taxonomy, and Evolution of Nonmarine Lineages within the American Grapsoid Crabs (Crustacea: Brachyura). Molecular Phylogenetics and Evolution. 15, 179–190. doi:10.1006/mpev.1999.0754.

Schubart, C.D., Neigel, J.E., and Felder, D.L. (2000b). The use of the mitochondrial 16S rRNA gene for phylogenetic and biogeographic studies of Crustacea. in Crustacean and the biodiversity crisis and Crustacea. Proceedings of the Fourth International Crustacean Congress, ed. F. R. Schram, J.C. von Vaupel Klein, Amsterdam, the Netherlands, 20-24 July 1998, vol. 2. (Leiden: Brill), 817–830.

Shahdadi, A., and Schubart, C. D. (2017). Taxonomic review of Perisesarma (Decapoda: Brachyura: Sesarmidae) and closely related genera based on morphology and molecular phylogenetics: new classification, two new genera and the questionable phylogenetic value of the epibranchial tooth. Zoological Journal of the Linnean Society. 182, 517–548. doi:10.1093/zoolinnean/zlx032.

Shahdadi, A., and Schubart, C.D. (2015). Evaluating the consistency and taxonomic importance of cheliped and other morphological characters that potentially allow identification of species of the genus Perisesarma De Man, 1895 (Brachyura, Sesarmidae). Crustaceana. 88, 1079–1095. 1079e1095. https://pred.uni-regensburg.de/id/eprint/6142

Shahdadi, A., Davie, P. J. F., and Schubart, C. D. (2019). A new species of Parasesarma (Decapoda: Brachyura: Sesarmidae) from northern Australian mangroves and its distinction from morphologically similar species. Zoologischer Anzeiger. 279, 116–125. doi:10.1016/j.jcz.2019.01.005.

Shin, S., Jung, S., Heller, K., Menzel, F., Hong, T. K., Shin, J. S., Lee, S. H., Lee, H., & Lee, S. (2014). DNA barcoding ofBradysia(Diptera: Sciaridae) for detection of the immature stages on agricultural crops. Journal of Applied Entomology, 139(8), 638–645. https://doi.org/10.1111/jen.12198

Silva, I. C., Mesquita, N., and Paula, J. (2010). Genetic and morphological differentiation of the mangrove crab Perisesarma guttatum (Brachyura: Sesarmidae) along an East African latitudinal gradient. Biological Journal of the Linnean Society. 99, 28–46.

Smith, J. M., Smith, N. H., O’Rourke, M., and Spratt, B. G. (1993). How clonal are bacteria? Proceedings of the National Academy of Sciences. 90, 4384–4388. doi:10.1073/pnas.90.10.4384.

Spalding, M., Kainuma, M., Collins, L. (2010). World Atlas of Mangroves (version 3.0). A collaborative project of ITTO, ISME, FAO, UNEP-WCMC, UNESCO-MAB, UNU-INWEH and TNC. London (UK): Earthscan, London. URL: http://www.routledge.com/books/details/9781844076574; http://data.unepwcmc.org/datasets/5

Tan, C. G. S., and Ng, P. K. L. (1994). An annotated checklist of mangrove brachyuran crabs from Malaysia and Singapore. Hydrobiologia. 285, 75–84. doi:10.1007/bf00005655.

Thangaraj M, Annadurai D, Ramesh T, Kumaran R & Ravitchandirane V (2020) DNA barcoding and preliminary phylogenetic analysis of few gastropods (Family: Potamididae and Nassariidae) in Vellar estuary mangroves, India by COI and 18S rRNA genes. Ind. J of Geo. Mar. Sci. 49(4): 596–600.

Thirumaraiselvi, R., Das, S., Ramanadevi, V., & Thangaraj, M. (2015). MtDNA Barcode Identification of Finfish Larvae from Vellar Estuary, Tamilnadu, India. Notulae Scientia Biologicae, 7(1). https://doi.org/10.15835/nsb.7.1.9478

Van Lavieren, H., Spalding, M., Alongi, D., Kainuma, M., Clüsener-Godt, M., and Adeel, Z. (2012). Securing the Future of Mangroves. A Policy Brief. UNU-INWEH, UNESCO-MAB with ISME, ITTO, FAO, UNEP-WCMC and TNC, 53.

von Cräutlein, M., Korpelainen, H., Pietiläinen, M. et al. DNA barcoding: a tool for improved taxon identification and detection of species diversity. Biodivers Conserv 20, 373–389 (2011). https://doi.org/10.1007/s10531-010-9964-0

Weigand, H., Beermann, A. J., Čiampor, F., Costa, F. O., Csabai, Z., Duarte, S., et al. (2019). DNA barcode reference libraries for the monitoring of aquatic biota in Europe: Gap-analysis and recommendations for future work. Science of The Total Environment. 678, 499–524. doi:10.1016/j.scitotenv.2019.04.247.

