## Supplementary material for "Diversity, phylogeny and DNA barcoding of brachyuran crabs in artificially created mangrove environments": Figure S2

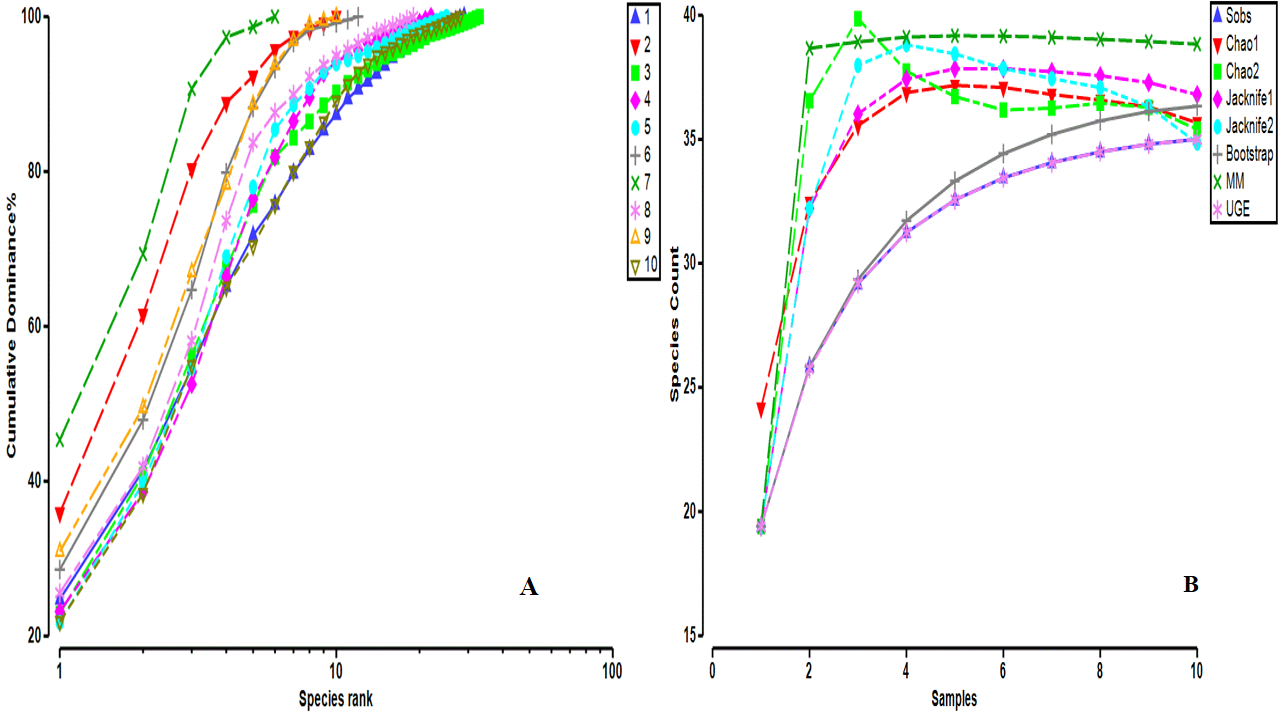

**Fig. S2:** Dominance plot (A) comparing the brachyuran crab diversity between different sampled stations. At stations, 3M and 7N, the dominance plot showed maximum and minimum diversity, respectively. Various species estimators (B) estimated the probability of obtaining 40 species. Chao 2 estimatd the maximum number of 40 in 3M (from which 33 species were sampled; 82.5% recovery) and UGE estimated a minimum of 35 species in 1A (from which 29 species were sampled; ~83% recovered).
