## Supplementary material for "Diversity, phylogeny and DNA barcoding of brachyuran crabs in artificially created mangrove environments": Figure S3

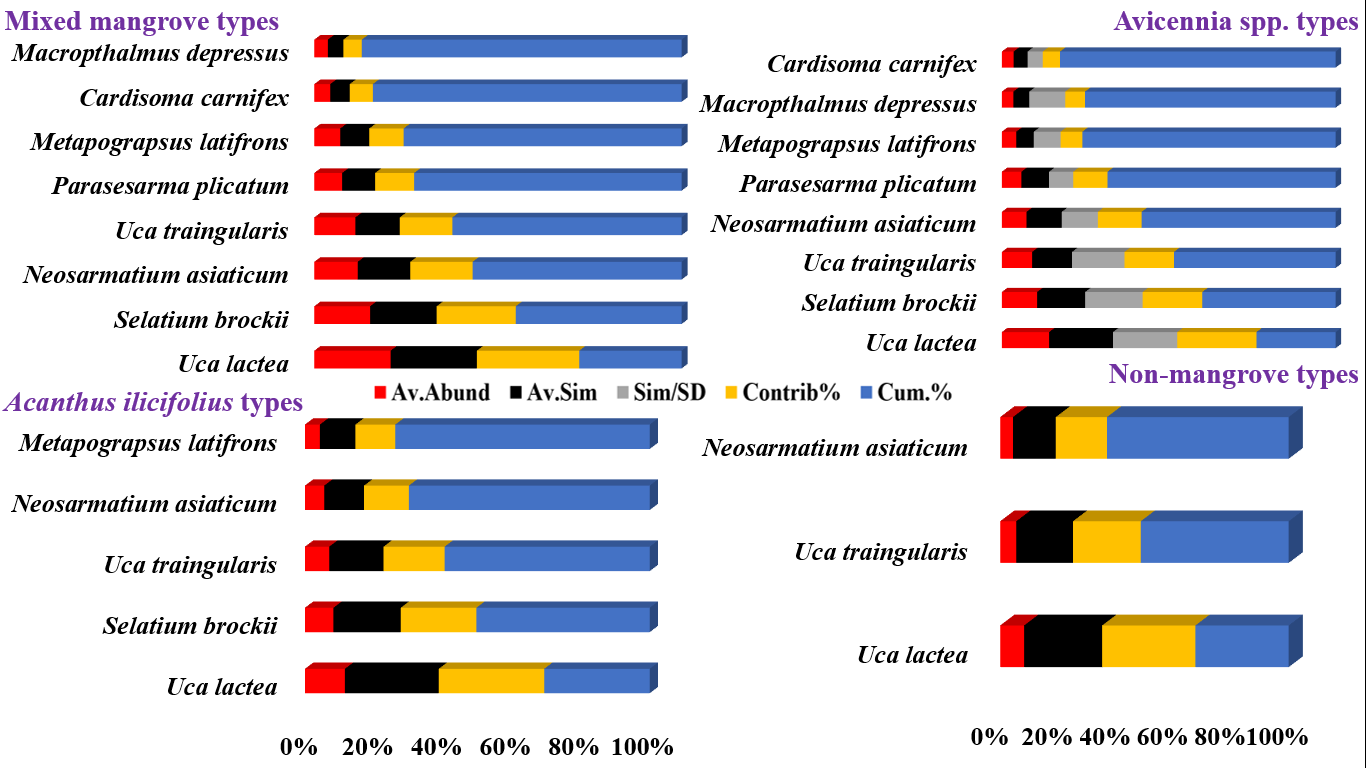

**Fig. S3:** The plot shows the patterns of similarity and dissimilarity between the brachyuran crab species composition and different mangrove types of sampled stations. Av.Abund- average abundances, Av.Sim- average similarity values, Sim/SD- similarity/ standard deviation, Contrib%- percentage of contribution, Cum.%- cumulative percentage. In all mangrove stations, *U. lactae* contributed average maximum similarity values.
