## Supplementary table 1 for "Diversity, phylogeny and DNA barcoding of brachyuran crabs in artificially created mangrove environments"

**Table S1.** Station-wise distribution of brachyuran crabs (average number/5m^2^).

| **Sl.No.** | **Name of the crab species** | **Family** | **Stations with type of zone** | | | | | | | | | |  | |
| --- | --- | --- | --- | --- | --- | --- | --- | --- | --- | --- | --- | --- | --- | --- |
|  |  |  | **1A** | **2N** | **3M** | **4A** | **5M** | **6Ai** | **7N** | **8A** | **9Ai** | **10A** | **Total** | **Percen**  **tage (%)** |
| **1** | ***Selatium brockii*** | Sesarmidae | 52 | 4 | 74 | 40 | 68 | 40 | 2 | 42 | 36 | 62 | **420** | **14.768** |
| **2** | ***Parasesarma plicatum*** | Sesarmidae | 26 | 1 | 30 | 26 | 34 | 12 | 0 | 26 | 10 | 20 | **185** | **6.505** |
| **3** | ***Perisesarma bidens*** | Sesarmidae | 0 | 0 | 1 | 1 | 1 | 0 | 0 | 0 | 0 | 1 | **4** | **0.141** |
| **4** | ***Neosarmatium asiaticum*** | Sesarmidae | 42 | 22 | 54 | 36 | 56 | 36 | 32 | 40 | 20 | 38 | **376** | **13.221** |
| **5** | ***Episesarma versicolor*** | Sesarmidae | 16 | 1 | 10 | 2 | 2 | 0 | 0 | 1 | 1 | 4 | **37** | **1.301** |
| **6** | ***E. mederi*** | Sesarmidae | 4 | 0 | 4 | 1 | 0 | 0 | 0 | 0 | 0 | 2 | **11** | **0.387** |
| **7** | ***Muradium tetragonum*** | Sesarmidae | 3 | 0 | 6 | 2 | 2 | 0 | 0 | 0 | 0 | 1 | **14** | **0.492** |
| **8** | ***Nanosesarma andersonii*** | Sesarmidae | 0 | 0 | 0 | 1 | 0 | 0 | 0 | 0 | 0 | 0 | **1** | **0.035** |
| **9** | ***N. minutum*** | Sesarmidae | 2 | 0 | 4 | 1 | 4 | 1 | 0 | 0 | 0 | 2 | **14** | **0.492** |
| **10** | ***N. batavicum*** | Sesarmidae | 1 | 0 | 2 | 0 | 1 | 0 | 0 | 0 | 0 | 0 | **4** | **0.141** |
| **11** | ***Metopograpsus latifrons*** | Grapsidae | 16 | 4 | 39 | 12 | 28 | 20 | 2 | 6 | 22 | 20 | **169** | **5.942** |
| **12** | ***M. frontalis*** | Grapsidae | 2 | 0 | 2 | 2 | 2 | 1 | 0 | 2 | 0 | 4 | **15** | **0.527** |
| **13** | ***Grapsus albolineatus*** | Grapsidae | 1 | 0 | 1 | 0 | 0 | 0 | 0 | 0 | 0 | 0 | **2** | **0.070** |
| **14** | ***G. tenuicrustatus*** | Grapsidae | 4 | 1 | 2 | 4 | 2 | 1 | 0 | 6 | 0 | 10 | **30** | **1.055** |
| **15** | ***Cardisoma carnifex*** | Gecarcinidae | 8 | 10 | 12 | 14 | 12 | 4 | 10 | 4 | 6 | 16 | **96** | **3.376** |
| **16** | ***Macrophthalmus depressus*** | Macrophthalmidae | 10 | 0 | 10 | 8 | 8 | 8 | 0 | 10 | 4 | 12 | **70** | **2.461** |
| **17** | ***M. erato*** | Macrophthalmidae | 8 | 0 | 1 | 1 | 0 | 0 | 0 | 0 | 0 | 0 | **10** | **0.352** |
| **18** | ***Ptychognathus altimanus*** | Varunidae | 1 | 0 | 2 | 1 | 1 | 0 | 0 | 1 | 0 | 0 | **6** | **0.211** |
| **19** | ***Pseudograpsus intermedius*** | Varunidae | 3 | 0 | 1 | 1 | 2 | 0 | 0 | 2 | 0 | 1 | **10** | **0.352** |
| **20** | ***Uca lactea*** | Ocypodidae | 98 | 42 | 110 | 60 | 82 | 68 | 68 | 66 | 60 | 82 | **736** | **25.879** |
| **21** | ***U. triangularis*** | Ocypodidae | 66 | 30 | 86 | 36 | 54 | 46 | 36 | 42 | 34 | 62 | **492** | **17.300** |
| **22** | ***Ocypode platytarsis*** | Ocypodidae | 1 | 0 | 2 | 0 | 1 | 0 | 0 | 0 | 0 | 8 | **12** | **0.422** |
| **23** | ***O. macrocera*** | Ocypodidae | 4 | 0 | 1 | 0 | 2 | 0 | 0 | 2 | 0 | 6 | **15** | **0.527** |
| **24** | ***O. brevicornis*** | Ocypodidae | 1 | 0 | 0 | 0 | 0 | 0 | 0 | 0 | 0 | 1 | **2** | **0.070** |
| **25** | ***Dotilla myctiroides*** | Dotillidae | 12 | 2 | 1 | 8 | 8 | 0 | 0 | 2 | 0 | 12 | **45** | **1.582** |
| **26** | ***D. intermedia*** | Dotillidae | 2 | 0 | 4 | 0 | 1 | 0 | 0 | 1 | 0 | 2 | **10** | **0.352** |
| **27** | ***Scylla serrata*** | Portunidae | 5 | 0 | 8 | 1 | 2 | 1 | 0 | 3 | 1 | 2 | **23** | **0.809** |
| **28** | ***S. olivacea*** | Portunidae | 4 | 0 | 2 | 0 | 0 | 0 | 0 | 1 | 0 | 1 | **8** | **0.281** |
| **29** | ***Portunus pelagicus*** | Portunidae | 0 | 0 | 2 | 0 | 0 | 0 | 0 | 0 | 0 | 1 | **3** | **0.105** |
| **30** | ***P. sanguinolentus*** | Portunidae | 0 | 0 | 1 | 0 | 0 | 0 | 0 | 0 | 0 | 2 | **3** | **0.105** |
| **31** | ***P. reticulatus*** | Portunidae | 1 | 0 | 4 | 1 | 0 | 0 | 0 | 0 | 0 | 1 | **7** | **0.246** |
| **32** | ***Charybdis feriata*** | Portunidae | 1 | 0 | 1 | 0 | 1 | 0 | 0 | 0 | 0 | 1 | **4** | **0.141** |
| **33** | ***C. lucifera*** | Portunidae | 0 | 0 | 1 | 0 | 0 | 0 | 0 | 0 | 0 | 0 | **1** | **0.035** |
| **34** | ***Thalamita chaptalii*** | Portunidae | 0 | 0 | 1 | 0 | 1 | 0 | 0 | 0 | 0 | 0 | **2** | **0.070** |
| **35** | ***T. crenata*** | Portunidae | 2 | 0 | 1 | 0 | 2 | 0 | 0 | 1 | 0 | 1 | **7** | **0.246** |
|  | **Total** |  | **396** | **117** | **480** | **259** | **377** | **238** | **150** | **258** | **194** | **375** | **2844** | **1 100.00** |

**A- *Avicennia* zone, N- Non-mangrove zone, M - Mixed mangrove zone, Ai - *Acanthus ilicifolius* zone**
