## Supplementary table 2 for "Diversity, phylogeny and DNA barcoding of brachyuran crabs in artificially created mangrove environments"

|  | | 1A | | 2N | | 3M | | | 4A | | 5M | 6Ai | 7N | | 8A | 9Ai | 10A |
| --- | --- | --- | --- | --- | --- | --- | --- | --- | --- | --- | --- | --- | --- | --- | --- | --- | --- |
| **Diversity indices** | | | | | | | | | | | | | | | | | |
| S | | 29 | | 10 | | **33** | | | 22 | | 25 | 12 | **6** | | 19 | 10 | 28 |
| N | | 396 | | **117** | | **480** | | | 259 | | 377 | 238 | 150 | | 258 | 194 | 375 |
| d | | 6.3 | | 2.69 | | **7** | | | 5.14 | | 5.59 | 2.9 | **1.5** | | 4.5 | 2.5 | 6.2 |
| J' | | **0.9** | | 0.9 | | 0.9 | | | 0.9 | | 0.89 | 0.9 | **0.9** | | 0.9 | 0.9 | 0.9 |
| Brillouin | | 2.7 | | 1.73 | | **2.8** | | | 2.37 | | 2.53 | 1.9 | **1.4** | | 2.3 | 1.8 | 2.6 |
| Fisher | | 15.8 | | 5.52 | | **18.5** | | | 12.66 | | 13.47 | 5.4 | **2.4** | | 10.3 | 4.4 | 15.4 |
| ES | -25 | **15.7** | | 9.55 | | 15.3 | | | 13.94 | | 13.55 | 9.9 | **5.9** | | 12.6 | 9 | 14.9 |
|  | -50 | 23.2 | | 10 | | **23.5** | | | 20.14 | | 20.08 | 12 | **6** | | 18.1 | 10 | 22.2 |
|  | -75 | 27.9 | | 10 | | **29.7** | | | 22 | | 25 | 12 | **6** | | 19 | 10 | 27.3 |
|  | -100 | 29 | | 10 | | **33** | | | 22 | | 25 | 12 | **6** | | 19 | 10 | 28 |
|  | -150 | 29 | | 10 | | **33** | | | 22 | | 25 | 12 | **6** | | 19 | 10 | 28 |
| H' | (log2) | 4.5 | | 3 | | **4.5** | | | 4.05 | | 4.16 | 3.2 | **2.3** | | 3.8 | 3.1 | 4.4 |
| Lambda' | | **0.04** | | 0.11 | | 0.04 | | | 0.05 | | 0.05 | 0.1 | **0.2** | | 0.06 | 0.1 | 0.04 |
| 1-Lambda' | | **0.9** | | 0.88 | | 0.9 | | | 0.94 | | 0.94 | 0.9 | **0.8** | | 0.9 | 0.9 | 0.9 |
| Delta* | | 93 | | **90.5** | | 92.8 | | | 92.59 | | 92.68 | 91.3 | 86.3 | | **93.3** | 91.4 | 93.1 |
| sDelta+ | | 2738.1 | | 933.33 | | **3089.6** | | | 2047.61 | | 2358.33 | 1127.3 | **560** | | 1822.2 | 940.7 | 2617.3 |
| sPhi+ | | 1866.7 | | 800 | | **2033.3** | | | 1566.66 | | 1733.33 | 933.3 | **500** | | 1400 | 833.3 | 1833.3 |
| **Species estimators** | | | | | | | | | | | | | | | | | |
| Sobs | | | 19.44 | | 25.83 | | 29.17 | 31.24 | | 32.55 | | 33.4 | 34.1 | 34.5 | | 34.8 | 35 |
| Sobs(SD) | | | 8.94 | | 6.87 | | 4.98 | 3.45 | | 2.33 | | 1.56 | 1.1 | 0.7 | | 0.4 | 0 |
| Chao1 | | | 24.17 | | 32.45 | | 35.54 | 36.88 | | 37.17 | | 37.0 | 36.8 | 36.6 | | 36.3 | 35.7 |
| Chao1(SD) | | | 4.7 | | 6.59 | | 6.574 | 6.17 | | 5.32 | | 4.5 | 3.6 | 2.9 | | 2.3 | 1.3 |
| Chao2 | | | 19.4 | | 36.55 | | **39.86** | 37.74 | | 36.71 | | 36.2 | 36.2 | 36.4 | | 36.3 | 35.4 |
| Chao2(SD) | | | 4.7 | | 7.09 | | 8.18 | 5.47 | | 3.92 | | 2.9 | 2.6 | 2.5 | | 2.1 | 0.9 |
| Jacknife1 | | | 19.56 | | 32.21 | | 36.01 | 37.42 | | 37.83 | | 37.8 | 37.7 | 37.6 | | 37.3 | 36.8 |
| Jacknife2 | | | 19.6 | | 32.21 | | 37.97 | 38.8 | | 38.44 | | 37.8 | 37.4 | 37.1 | | 36.3 | 34.8 |
| Bootstrap | | | 19.5 | | 25.85 | | 29.35 | 31.71 | | 33.31 | | 34.4 | 35.2 | 35.8 | | 36.1 | 36.3 |
| MM | | | 19.4 | | 38.66 | | 38.92 | 39.11 | | 39.17 | | 39.1 | 39.1 | 39 | | 38.9 | 38.8 |
| UGE | | | 19.4 | | 25.75 | | 29.18 | 31.24 | | 32.55 | | 33.4 | 34 | 34.5 | | 34.8 | **35** |

Table S2: Table representing the values of the various diversity indices and species estimators. Values marked in bold represents higher or lower values among the parameters in question.

S - Number of species; N - Abundance (number of individuals); d - Margalef index; J’- Pielou’s evenness index; Brillouin index; Fisher’s alpha index; Expected number of species ES(25); ES(50); ES(75); ES(100); ES(150); H'(log2) – Shannon -Wiener diversity index; Lambda'- Simpson’s dominance index; 1-Lambda' - Simpson richness index; Delta - Taxonomic diversity; sDelta+ - Total Taxonomic distinctness index and sPhi+ - Total Phylogenetic diversity index. MM- Michaelis Menton; UGE- Ugland-Gray-Ellingsen.
