## Supplementary table 3 for "Diversity, phylogeny and DNA barcoding of brachyuran crabs in artificially created mangrove environments"

**Table S4**. Index of association values between various brachyuran crab species was shown. *Uca* spp. exhibited maximum index values (97.7%) whereas the minimum association values (0%) were observed between species such as *Nanosesarma batavicum*, *Grapsus albolineatus*, *Ocypode platytarsis*, *O. macrocera*, *O. brevicornis*, *Dotilla intermedia*, *Scylla olivacea*, *Portunus pelagicus*, *P. sanguinolentus*, *Charybdis feriatus*, *C. lucifera*, *Thalamita chaptalii* and *T. crenata* with *N. andersonii* and *C. lucifera.* *T. chaptalii* also had minimum association value (0%) with *O. brevicornis*.

|  |  | **1** | **2** | **3** | **4** | **5** | **6** | **7** | **8** | **9** | **10** | **11** | **12** | **13** | **14** | **15** | **16** | **17** | **18** | **19** | **20** | **21** | **22** | **23** | **24** | **25** | **26** | **27** | **28** | **29** | **30** | **31** | **32** | **33** | **34** | **35** |
| --- | --- | --- | --- | --- | --- | --- | --- | --- | --- | --- | --- | --- | --- | --- | --- | --- | --- | --- | --- | --- | --- | --- | --- | --- | --- | --- | --- | --- | --- | --- | --- | --- | --- | --- | --- | --- |
| **1** | ***Selatium brockii*** |  |  |  |  |  |  |  |  |  |  |  |  |  |  |  |  |  |  |  |  |  |  |  |  |  |  |  |  |  |  |  |  |  |  |  |
| **2** | ***Parasesarma plicatum*** | **92.1** |  |  |  |  |  |  |  |  |  |  |  |  |  |  |  |  |  |  |  |  |  |  |  |  |  |  |  |  |  |  |  |  |  |  |
| **3** | ***Perisesarma bidens*** | **51.3** | **53.9** |  |  |  |  |  |  |  |  |  |  |  |  |  |  |  |  |  |  |  |  |  |  |  |  |  |  |  |  |  |  |  |  |  |
| **4** | ***Neosarmatium asiaticum*** | **88.6** | **84.6** | **44.5** |  |  |  |  |  |  |  |  |  |  |  |  |  |  |  |  |  |  |  |  |  |  |  |  |  |  |  |  |  |  |  |  |
| **5** | ***Episesarma versicolor*** | **74.7** | **73.7** | **53.3** | **71.9** |  |  |  |  |  |  |  |  |  |  |  |  |  |  |  |  |  |  |  |  |  |  |  |  |  |  |  |  |  |  |  |
| **6** | ***Episesarma mederi*** | **49.6** | **52.1** | **62.6** | **42.9** | **70.6** |  |  |  |  |  |  |  |  |  |  |  |  |  |  |  |  |  |  |  |  |  |  |  |  |  |  |  |  |  |  |
| **7** | ***Muradium tetragonum*** | **62.7** | **67.1** | **72.8** | **55.2** | **74.1** | **80.3** |  |  |  |  |  |  |  |  |  |  |  |  |  |  |  |  |  |  |  |  |  |  |  |  |  |  |  |  |  |
| **8** | ***Nanosesarma andersonii*** | **10.5** | **13.2** | **25.0** | **9.9** | **9.4** | **15.6** | **17.7** |  |  |  |  |  |  |  |  |  |  |  |  |  |  |  |  |  |  |  |  |  |  |  |  |  |  |  |  |
| **9** | ***Nanosesarma minutum*** | **73.7** | **74.2** | **72.7** | **65.1** | **69.3** | **66.0** | **80.1** | **11.3** |  |  |  |  |  |  |  |  |  |  |  |  |  |  |  |  |  |  |  |  |  |  |  |  |  |  |  |
| **10** | ***Nanosesarma batavicum*** | **39.8** | **42.4** | **50.0** | **35.2** | **57.2** | **60.5** | **69.9** | **0.0** | **61.3** |  |  |  |  |  |  |  |  |  |  |  |  |  |  |  |  |  |  |  |  |  |  |  |  |  |  |
| **11** | ***Metapograpsus latifrons*** | **91.2** | **84.9** | **50.6** | **86.5** | **74.9** | **47.2** | **61.0** | **9.0** | **72.3** | **40.4** |  |  |  |  |  |  |  |  |  |  |  |  |  |  |  |  |  |  |  |  |  |  |  |  |  |
| **12** | ***Metapograpsus frontalis*** | **83.7** | **88.1** | **62.0** | **75.6** | **67.0** | **62.0** | **68.7** | **14.0** | **79.4** | **42.1** | **75.1** |  |  |  |  |  |  |  |  |  |  |  |  |  |  |  |  |  |  |  |  |  |  |  |  |
| **13** | ***Grapsus albolineatus*** | **26.1** | **27.3** | **25.0** | **22.8** | **47.8** | **62.4** | **52.2** | **0.0** | **38.7** | **70.7** | **26.6** | **28.1** |  |  |  |  |  |  |  |  |  |  |  |  |  |  |  |  |  |  |  |  |  |  |  |
| **14** | ***Grapsus tenuicrustatus*** | **75.9** | **80.2** | **55.3** | **74.6** | **69.2** | **59.4** | **59.8** | **13.9** | **67.7** | **33.4** | **69.1** | **88.1** | **23.6** |  |  |  |  |  |  |  |  |  |  |  |  |  |  |  |  |  |  |  |  |  |  |
| **15** | ***Cardisoma carnifex*** | **82.7** | **80.0** | **48.5** | **90.0** | **72.8** | **46.4** | **57.1** | **12.4** | **63.4** | **32.2** | **82.8** | **71.0** | **20.8** | **74.6** |  |  |  |  |  |  |  |  |  |  |  |  |  |  |  |  |  |  |  |  |  |
| **16** | ***Macropthalmus depressus*** | **90.7** | **92.7** | **52.4** | **82.7** | **72.5** | **53.8** | **63.6** | **12.1** | **76.5** | **39.1** | **83.1** | **89.3** | **27.0** | **80.3** | **78.8** |  |  |  |  |  |  |  |  |  |  |  |  |  |  |  |  |  |  |  |  |
| **17** | ***Macropthalmus erato*** | **36.6** | **40.5** | **41.4** | **32.7** | **56.8** | **67.5** | **60.0** | **20.7** | **48.1** | **50.0** | **35.6** | **42.1** | **70.7** | **37.5** | **33.1** | **39.1** |  |  |  |  |  |  |  |  |  |  |  |  |  |  |  |  |  |  |  |
| **18** | ***Ptychognathus altimanus*** | **61.0** | **68.7** | **61.9** | **55.5** | **65.1** | **60.2** | **79.9** | **18.5** | **68.5** | **63.1** | **55.7** | **70.2** | **44.6** | **64.3** | **51.2** | **64.6** | **57.7** |  |  |  |  |  |  |  |  |  |  |  |  |  |  |  |  |  |  |
| **19** | ***Pseudograpsus intermedius*** | **73.0** | **79.4** | **58.4** | **65.7** | **74.9** | **62.6** | **78.2** | **13.2** | **72.5** | **54.8** | **64.3** | **81.8** | **36.1** | **76.9** | **64.4** | **77.6** | **49.4** | **81.9** |  |  |  |  |  |  |  |  |  |  |  |  |  |  |  |  |  |
| **20** | ***Uca lactea*** | **88.3** | **83.6** | **42.7** | **96.3** | **73.2** | **43.7** | **54.3** | **9.1** | **64.0** | **34.6** | **87.0** | **73.6** | **24.0** | **74.4** | **89.8** | **82.1** | **33.1** | **53.3** | **63.9** |  |  |  |  |  |  |  |  |  |  |  |  |  |  |  |  |
| **21** | ***Uca traingularis*** | **89.1** | **84.9** | **44.1** | **95.2** | **74.6** | **45.2** | **55.8** | **8.7** | **65.6** | **35.8** | **87.9** | **75.0** | **25.1** | **74.6** | **89.4** | **83.4** | **33.8** | **53.8** | **65.0** | **97.7** |  |  |  |  |  |  |  |  |  |  |  |  |  |  |  |
| **22** | ***Ocypode platytarsis*** | **52.8** | **53.9** | **63.7** | **45.3** | **59.9** | **60.7** | **67.2** | **0.0** | **70.7** | **54.7** | **52.0** | **62.0** | **38.7** | **55.3** | **45.4** | **53.8** | **36.7** | **54.7** | **58.5** | **45.2** | **47.1** |  |  |  |  |  |  |  |  |  |  |  |  |  |  |
| **23** | ***Ocypode macrocera*** | **61.4** | **65.0** | **54.2** | **55.7** | **65.7** | **58.3** | **63.3** | **0.0** | **61.2** | **53.3** | **54.2** | **74.1** | **36.2** | **72.3** | **52.0** | **65.9** | **36.2** | **64.7** | **82.4** | **54.5** | **55.2** | **73.7** |  |  |  |  |  |  |  |  |  |  |  |  |  |
| **24** | ***Ocypode brevicornis*** | **24.9** | **24.7** | **25.0** | **20.9** | **40.0** | **53.2** | **34.1** | **0.0** | **32.0** | **29.3** | **22.0** | **33.9** | **50.0** | **35.7** | **22.6** | **28.3** | **50.0** | **18.5** | **36.1** | **22.3** | **23.1** | **61.3** | **53.8** |  |  |  |  |  |  |  |  |  |  |  |  |
| **25** | ***Dotilla myctiroides*** | **67.1** | **70.3** | **61.7** | **65.5** | **72.7** | **63.9** | **74.1** | **17.2** | **66.7** | **44.4** | **62.4** | **76.7** | **27.2** | **80.2** | **67.7** | **67.1** | **44.4** | **67.6** | **79.5** | **64.3** | **65.0** | **59.2** | **74.0** | **42.2** |  |  |  |  |  |  |  |  |  |  |  |
| **26** | ***Dotilla intermedia*** | **63.5** | **66.7** | **60.4** | **55.8** | **71.3** | **70.7** | **77.1** | **0.0** | **69.3** | **64.6** | **58.3** | **76.0** | **50.0** | **68.8** | **52.0** | **67.3** | **41.4** | **73.9** | **76.5** | **54.8** | **56.5** | **74.0** | **82.8** | **41.4** | **70.8** |  |  |  |  |  |  |  |  |  |  |
| **27** | ***Scylla serrata*** | **83.0** | **86.7** | **52.7** | **77.8** | **80.7** | **59.2** | **70.4** | **7.9** | **76.7** | **51.3** | **79.1** | **80.0** | **40.1** | **73.2** | **72.2** | **86.6** | **46.3** | **73.0** | **75.0** | **78.6** | **80.1** | **60.8** | **65.9** | **28.9** | **62.7** | **76.2** |  |  |  |  |  |  |  |  |  |
| **28** | ***Scylla olivacea*** | **49.9** | **52.1** | **43.5** | **43.4** | **67.8** | **75.8** | **60.2** | **0.0** | **54.7** | **55.4** | **44.6** | **60.6** | **63.1** | **59.1** | **40.6** | **55.3** | **57.7** | **63.1** | **67.8** | **44.2** | **45.9** | **57.1** | **71.8** | **55.4** | **54.3** | **79.9** | **65.0** |  |  |  |  |  |  |  |  |
| **29** | ***Portunus pelagicus*** | **27.2** | **25.7** | **50.0** | **22.3** | **34.4** | **53.2** | **43.1** | **0.0** | **38.7** | **41.4** | **27.8** | **33.9** | **50.0** | **31.7** | **24.7** | **28.3** | **20.7** | **26.1** | **26.5** | **23.0** | **24.8** | **64.1** | **41.7** | **41.4** | **27.2** | **50.0** | **33.6** | **44.6** |  |  |  |  |  |  |  |
| **30** | ***Portunus sanguinolentus*** | **27.2** | **25.7** | **50.0** | **22.3** | **34.4** | **53.2** | **43.1** | **0.0** | **38.7** | **41.4** | **27.8** | **33.9** | **41.4** | **31.7** | **24.7** | **28.3** | **20.7** | **26.1** | **26.5** | **23.0** | **24.8** | **68.0** | **41.7** | **50.0** | **27.2** | **50.0** | **33.6** | **44.6** | **82.8** |  |  |  |  |  |  |
| **31** | ***Portunus reticulatus*** | **49.6** | **52.1** | **65.0** | **42.9** | **63.9** | **86.8** | **80.7** | **20.0** | **66.0** | **60.0** | **47.2** | **62.0** | **60.0** | **57.5** | **46.4** | **53.8** | **60.7** | **63.1** | **59.7** | **43.7** | **45.2** | **58.7** | **52.1** | **40.0** | **63.3** | **69.3** | **59.2** | **64.6** | **60.0** | **60.0** |  |  |  |  |  |
| **32** | ***Charybdis feriatus*** | **52.8** | **53.9** | **75.0** | **45.3** | **68.9** | **72.0** | **76.8** | **0.0** | **77.3** | **75.0** | **52.0** | **62.0** | **50.0** | **55.3** | **45.4** | **53.8** | **45.7** | **61.9** | **68.1** | **45.2** | **47.1** | **79.7** | **78.3** | **50.0** | **65.5** | **81.1** | **62.5** | **68.5** | **50.0** | **50.0** | **65.0** |  |  |  |  |
| **33** | ***Charybdis lucifera*** | **14.2** | **14.2** | **25.0** | **12.1** | **21.1** | **31.2** | **30.6** | **0.0** | **22.7** | **41.4** | **16.2** | **14.0** | **50.0** | **9.8** | **11.4** | **13.5** | **20.7** | **26.1** | **13.2** | **12.3** | **13.4** | **22.7** | **12.1** | **0.0** | **6.1** | **29.3** | **22.4** | **26.1** | **58.6** | **41.4** | **40.0** | **25.0** |  |  |  |
| **34** | ***Thalamita chaptalii*** | **27.9** | **29.2** | **50.0** | **24.5** | **30.5** | **31.2** | **48.2** | **0.0** | **45.3** | **70.7** | **30.0** | **28.1** | **50.0** | **19.6** | **22.9** | **25.6** | **20.7** | **44.6** | **31.9** | **23.0** | **24.0** | **38.7** | **29.2** | **0.0** | **23.3** | **43.9** | **33.6** | **26.1** | **50.0** | **41.4** | **40.0** | **50.0** | **50.0** |  |  |
| **35** | ***Thalamita crenata*** | **63.5** | **67.1** | **58.6** | **55.8** | **70.9** | **58.6** | **68.9** | **0.0** | **71.8** | **65.7** | **58.3** | **73.3** | **41.4** | **67.6** | **52.0** | **67.3** | **41.4** | **71.3** | **85.2** | **54.8** | **56.5** | **66.4** | **87.6** | **41.4** | **70.2** | **84.3** | **71.0** | **75.7** | **34.3** | **34.3** | **54.3** | **82.8** | **17.2** | **41.4** |  |
