## Supplementary table 4 for "Diversity, phylogeny and DNA barcoding of brachyuran crabs in artificially created mangrove environments"

Table S4: The COI-K2P distance analysis between the various brachyuran crab species

|  | **1** | **2** | **3** | **4** | **5** | **6** | **7** | **8** | **9** | **10** | **11** | **12** | **13** | **14** | **15** |
| --- | --- | --- | --- | --- | --- | --- | --- | --- | --- | --- | --- | --- | --- | --- | --- |
| **(1) *Perisesarma bidens*** |  |  |  |  |  |  |  |  |  |  |  |  |  |  |  |
| **(2) *Parasesarma plicatum*** | 0.07 |  |  |  |  |  |  |  |  |  |  |  |  |  |  |
| **(3) *Neosarmatium asiaticum*** | 0.14 | 0.12 |  |  |  |  |  |  |  |  |  |  |  |  |  |
| **(4) *Episesarma versicolor*** | 0.13 | 0.11 | 0.09 |  |  |  |  |  |  |  |  |  |  |  |  |
| **(5) *Nanosesarma minutum*** | 0.13 | 0.11 | 0.11 | 0.10 |  |  |  |  |  |  |  |  |  |  |  |
| **(6) *Macrophthalmus depressus*** | 0.17 | 0.18 | 0.18 | 0.19 | 0.18 |  |  |  |  |  |  |  |  |  |  |
| **(7) *Plagusia dentipes*** | 0.20 | 0.22 | 0.19 | 0.18 | 0.17 | 0.18 |  |  |  |  |  |  |  |  |  |
| **(8) *Uca lactea*** | 0.24 | 0.23 | 0.21 | 0.22 | 0.20 | 0.21 | 0.21 |  |  |  |  |  |  |  |  |
| **(9) *Uca triangularis*** | 0.24 | 0.22 | 0.21 | 0.22 | 0.20 | 0.22 | 0.21 | 0.01 |  |  |  |  |  |  |  |
| **(10) *Grapsus albolineatus*** | 0.24 | 0.24 | 0.20 | 0.19 | 0.21 | 0.22 | 0.21 | 0.21 | 0.21 |  |  |  |  |  |  |
| **(11) *Ocypode platytarsis*** | 0.22 | 0.22 | 0.21 | 0.21 | 0.22 | 0.20 | 0.21 | 0.21 | 0.21 | 0.23 |  |  |  |  |  |
| **(12) *Ocypode brevicornis*** | 0.23 | 0.22 | 0.24 | 0.23 | 0.24 | 0.21 | 0.24 | 0.20 | 0.20 | 0.22 | 0.19 |  |  |  |  |
| **(13) *Cardisoma carnifex*** | 0.22 | 0.21 | 0.21 | 0.22 | 0.22 | 0.20 | 0.20 | 0.21 | 0.21 | 0.25 | 0.20 | 0.23 |  |  |  |
| **(14) *Metopograpsus latifrons*** | 0.20 | 0.20 | 0.20 | 0.18 | 0.20 | 0.22 | 0.21 | 0.21 | 0.21 | 0.21 | 0.21 | 0.23 | 0.20 |  |  |
| **(15) *Metopograpsus frontalis*** | 0.20 | 0.20 | 0.19 | 0.18 | 0.20 | 0.21 | 0.20 | 0.21 | 0.21 | 0.20 | 0.20 | 0.24 | 0.20 | 0.00 |  |
