## Supplementary figures and images for "Diversity, phylogeny and DNA barcoding of brachyuran crabs in artificially created mangrove environments"

### Figure S1

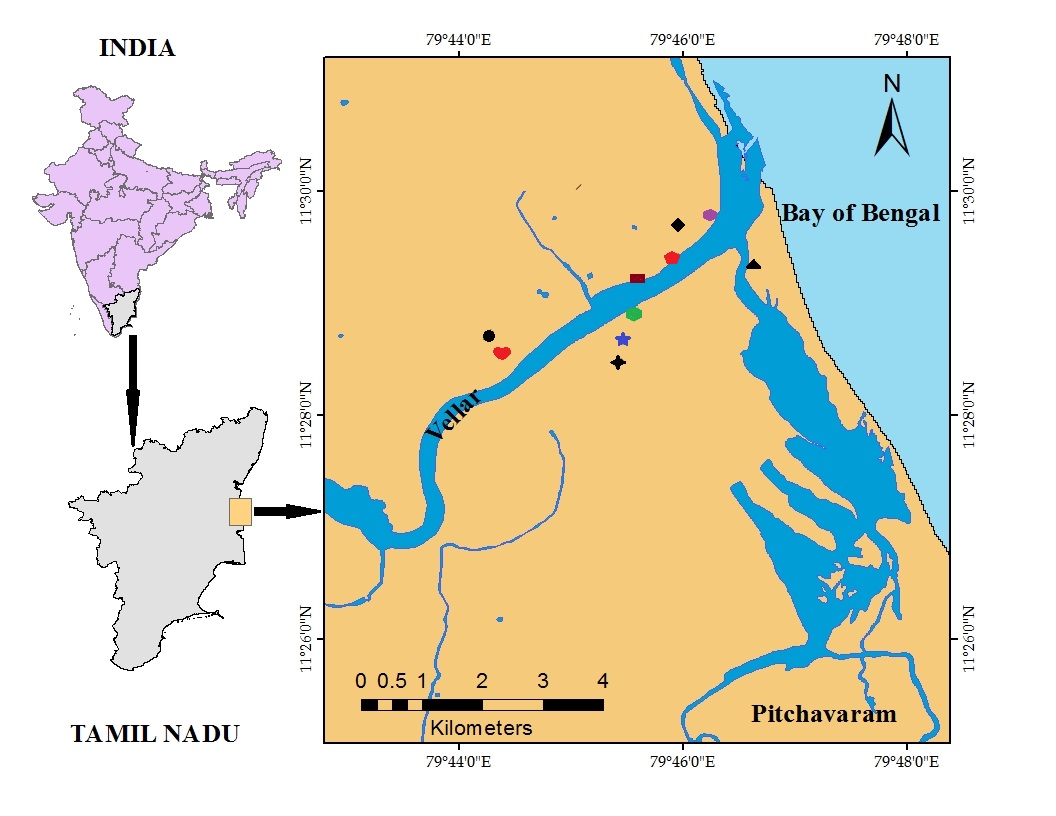
